# Cyclin-dependent kinase-like 5 (CDKL5) binds to talin and is anchored at the postsynaptic density via direct interaction with PDZ domains

**DOI:** 10.1101/2025.08.27.672337

**Authors:** Yasumi Otani, Emily Goodall, Vignesh Srinivasan, Neil J Ball, Samuel F. H. Barnett, Tobias Zech, Juha Saarikangas, Benjamin T. Goult

**Affiliations:** Department of Biochemistry, Cell & Systems Biology, Institute of Systems, Molecular & Integrative Biology, University of Liverpool, Crown Street, Liverpool L69 7ZB, U.K; Molecular & Clinical Cancer Medicine, Institute of Systems, Molecular & Integrative Biology, University of Liverpool, Crown Street, Liverpool L69 7ZB, U.K; Helsinki Institute of Life Science, HiLIFE, University of Helsinki, P.O. Box56, 00014, Helsinki, Finland; Faculty of Biological and Environmental Sciences, University of Helsinki, P.O. Box56, 00014, Helsinki, Finland; Genomics England, United Kingdom; Liverpool Interdisciplinary Neuroscience Centre, University of Liverpool, Liverpool, L9 7AL, U.K

## Abstract

Cyclin-dependent kinase-like 5 (CDKL5) is a serine/threonine kinase essential for brain development and function. Mutations in the *CDKL5* gene cause CDKL5 deficiency disorder (CDD), a severe early-onset epileptic encephalopathy characterised by defects in synapse formation and function. Despite extensive research, the molecular mechanisms by which CDKL5 mutations disrupt synaptic function and lead to epilepsy remain unclear. Here, we report that the major neuronal isoform of CDKL5 contains a C-terminal PDZ domain-binding motif. We demonstrate that this motif mediates interactions with the PDZ domains of PSD-95 and SHANK proteins, facilitating the recruitment of CDKL5 to the postsynaptic density. Disruption of CDKL5’s PDZ-binding motif results in its mislocalisation and impaired spine formation. Additionally, we show that CDKL5 directly interacts with the mechanosensitive synaptic scaffold protein talin, via the N-terminal kinase domain of CDKL5 and the R8 rod domain of talin. Our findings establish how CDKL5 is targeted to synapses and suggest that its activity may be spatially regulated through talin-mediated mechanical signalling. We propose that the spatial positioning of the CDKL5 kinase domain might be mechanically-operated and regulated by talin domain unfolding. As talin undergoes structural transitions in its force-dependent binary switch domains, the kinase domain bound to R8 would be moved up and down within the synaptic compartment as a function of the changing talin conformation. These insights enhance our understanding of the pathogenic mechanisms underlying CDKL5 variants with premature stop codons and highlight the need to re-evaluate studies that have used C-terminally tagged or the non-PDZ-binding isoform of CDKL5 to assess its neuronal function.

Graphical Abstract
Cyclin-dependent kinase-like 5 (CDKL5) is a mechanically regulated enzyme.
The C-terminal PDZ-binding motif in CDKL5 anchors CDKL5 at the postsynaptic density (PSD). **(A)** The large unstructured region serves to tether the CDKL5 kinase domain at the synapse. **i)** The kinase domain is restricted to the blue semi-circle, giving CDKL5 a zone of activity, with radius the length of the tether (>200 nm). **ii)** Epilepsy-causing mutations that introduce premature stop codons untether CDKL5. **(B)** The mechanosensitive synaptic scaffold protein talin is also tethered to the membrane by interaction with integrins and/or amyloid precursor protein (APP). The CDKL5-binding domain, R8 is shown in dark green. The force-dependent binary switch domains in the talin rod region open and close in response to force and change the length of the molecule and the location of the CDKL5-binding site. **(C)** As talin changes its conformation, the CDKL5-binding site on R8 is moved up and down within the synapse, as a result the kinase activity of CDKL5 is targeted to discrete locations within the synapse. Loss of this localisation or incorrect positioning of the kinase within the synapse would result in loss of synchronisation of the synapse in the context of the circuit it is part of and, we propose, result in re-entry circuits and epilepsy. See also [Supplementary Movie 1](https://www.youtube.com/watch?v=WC6t-A5GHPo).

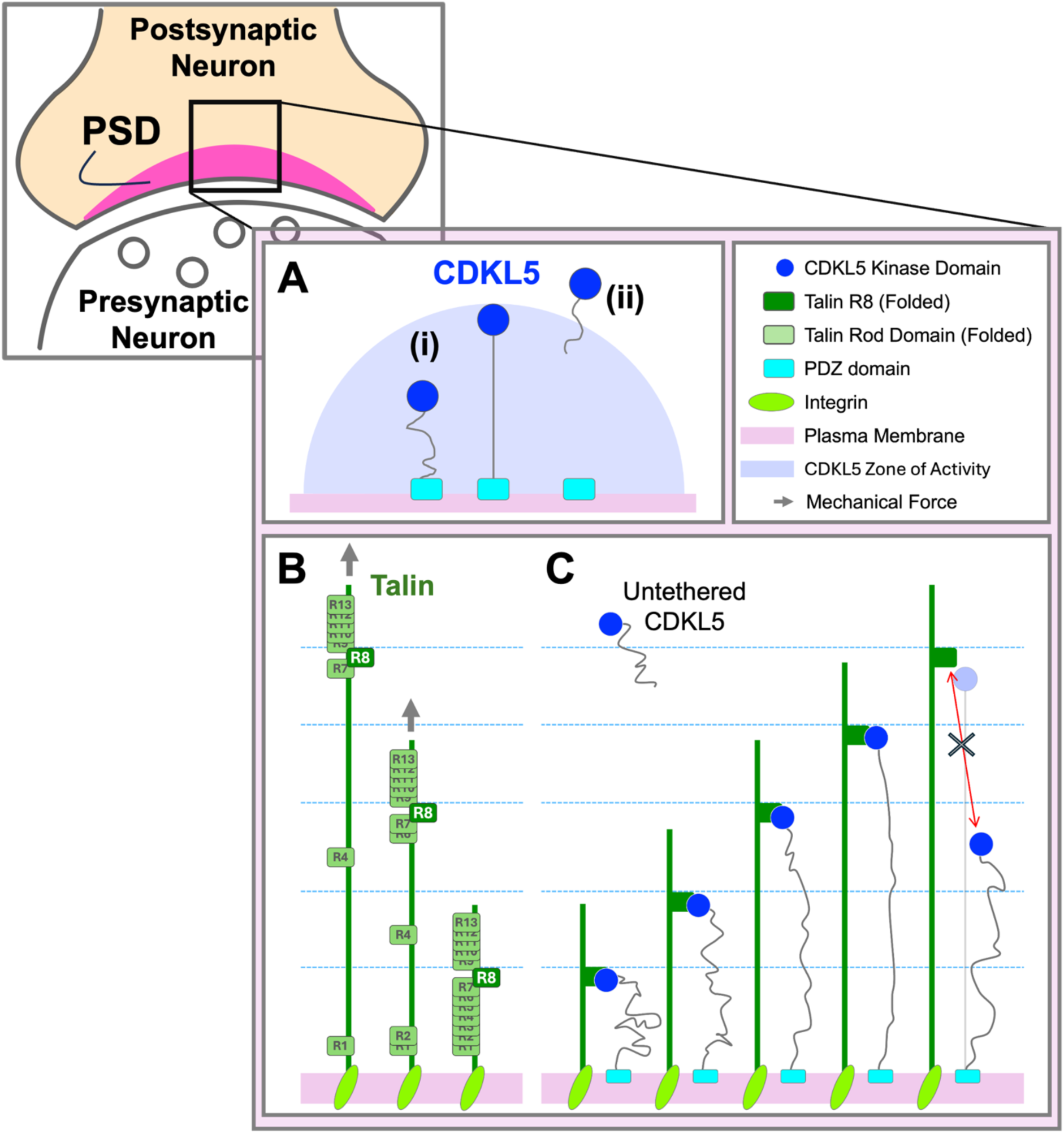

## Introduction

CDKL5 (Cyclin-dependent kinase-like 5) is a serine/threonine kinase (also known as serine/threonine-protein kinase 9, STK9) encoded by the X-linked *CDKL5* gene. CDKL5 plays multiple key roles in brain development and in neuronal processes (Kadam et al., 2019; Van Bergen et al., 2022). Loss-of-function mutations in CDKL5 result in two rare but devastating developmental disorders, CDKL5 deficiency disorder (CDD) and Rett Syndrome (RTT), which show similar phenotypes including epilepsy and developmental delays and mostly affect females (Kadam et al., 2019; Tao et al., 2004). CDD leads to severe early-onset epileptic encephalopathy, developmental and epileptic encephalopathy (DEE) and multiple CDD-causing variants of CDKL5 have been reported (You et al., 2023). RTT is associated with neurodevelopmental disorders causing mobile disabilities, mental-developmental delays and seizures, and CDKL5 variants were found in atypical RTT patients suffering from early drug-resistant epilepsy in addition to those typical RTT symptoms. 90-95% of RTT patients have mutations in another X-linked gene, *MECP2* encoding MeCP2 (methyl-CpG-binding protein 2), which is the cause of typical RTT (Kadam et al., 2019; Neul et al., 2008).

CDKL5 is a large protein of about 1000 residues, that is comprised of an N-terminal kinase domain, followed by a large unstructured C-terminal region (Fig. 1A). This large region extending beyond the kinase domain is the reason CDKL5 is classed as a CDK-like enzyme. There are two isoforms of CDKL5 which arise due to alternative splicing with isoform 1 being the predominantly expressed isoform in the brain (Williamson et al., 2012). The two isoforms differ only at the C-terminus (Fig. 1B). The kinase activity of CDKL5 has multiple roles in healthy brain function, including important roles in efficient synaptic vesicle endocytosis (Kontaxi et al., 2023) and in controlling synapse-connection strength and synaptic plasticity at postsynaptic sites (Okuda et al., 2017; Ricciardi et al., 2012). One example of a CDKL5 substrate is the synaptic adhesion molecule NGL-1 (Netrin-G ligand-1), and NGL-1 phosphorylation by CDKL5 has been shown to strengthen the association between NGL-1 and the postsynaptic density (PSD) scaffolding protein, PSD-95 (Postsynaptic density protein 95) (Ricciardi et al., 2012).

**Figure 1.**
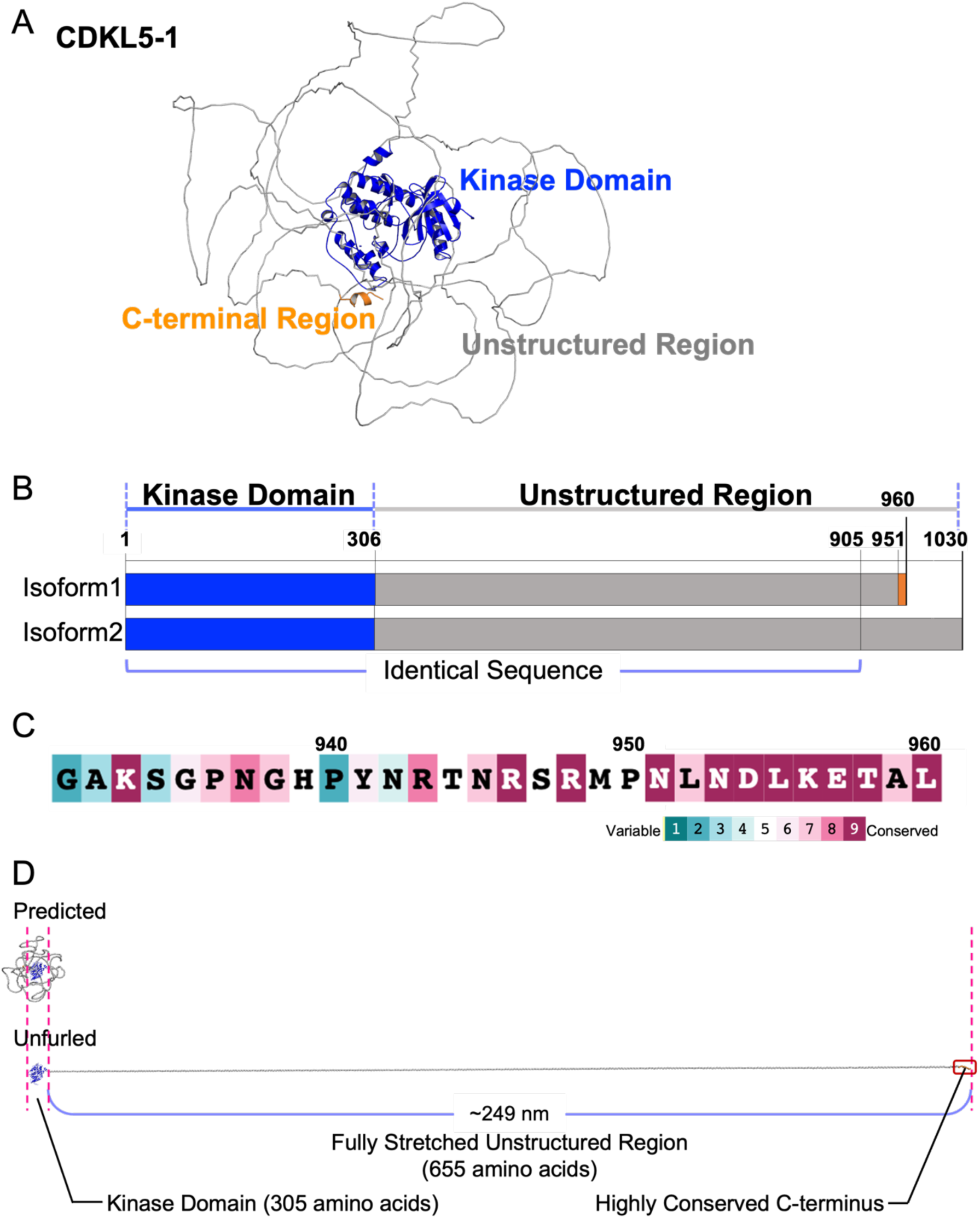
The neuronal CDKL5 isoform contains a highly conserved motif at the C-terminus. **(A)** Structural model of the neuronal CDKL5 isoform (CDKL5-1) protein (Human, UniProt ID: O76039-2) predicted by AlphaFold (Jumper et al., 2021). The kinase domain (blue), the large unstructured region (grey) and the conserved C-terminal motif (orange) are labelled. **(B)** The two CDKL5 isoforms, CDKL5-1 and CDKL5-2 vary only at the C-terminus. Both CDKL5 isoforms are identical in residues 1-905 which includes the kinase domain and the first 599 residues of the unstructured region. **(C)** The C-terminus of the neuronal CDKL5 isoform is highly conserved. The last 30 residues of CDKL5-1 (residues 931-960) are shown, coloured based on ConSurf conservation score (Yariv et al., 2023) where magenta denotes high conservation. **(D)** The unstructured region of CDKL5 is 654 amino acids long and able to stretch >200 nm when fully extended (see [Supplementary Movie 1](https://www.youtube.com/watch?v=WC6t-A5GHPo)). Top: The structural model of human CDKL5-1 as in (A). Bottom: The unfurled model of CDKL5-1 showing the dimensions of the protein and the length of the unstructured region.

### PDZ domains at the synapse

A common modular binding domain at the synapse is the PDZ (PSD-95/Discs-large/ZO-1) domain. PDZ domains comprise about 90 residues and are known as protein-protein interaction modules, typically binding to the very C-terminus of proteins. PSD-95 comprises three PDZ domains, an SH3 (Src homology 3) domain and a GK (guanylate kinase-like) domain (Fig. 3A), and plays a central role in organising the signalling machinery at the post-synaptic side of the synapse (Cho et al., 1992; Feng and Zhang, 2009; Hunt et al., 1996). The PDZ domains of PSD-95 proteins bind to the C-terminus of many proteins and are classified as class I PDZ domains based on their preferred binding motif (Ser/Thr-X-Φ-COOH, where X can be any residue and Φ is a hydrophobic amino acid) (Rodzli et al., 2020; Songyang et al., 1997). SHANK (SH3 and multiple ankyrin repeat domains) proteins are another class of PDZ domain containing proteins at the PSD (Sala et al., 2015; Tu et al., 1999). There are three SHANK family members, encoded by *Shank1*, *Shank2*, and *Shank3* genes, each containing a Shank/ProSAP N-terminal (SPN) domain, multiple ankyrin repeats, an SH3 domain, a class 1 PDZ domain, a proline rich region and a SAM (sterile α motif) domain at the C-terminus (Fig. 2A). Numerous SHANK isoforms exist due to multiple promoters and alternative splicing (Monteiro and Feng, 2017; Wang et al., 2014) and mutations in SHANK family genes are strongly linked to autism spectrum disorders (ASD) and other neurodevelopmental disorders (Boccuto et al., 2012; Jiang and Ehlers, 2013; Leblond et al., 2014; Sala et al., 2015). The PDZ domains of SHANK proteins have been shown to bind GKAP1 (also known as SAPAP1; SAP90/PSD-95-associated protein 1) forming a PSD-95/GKAP1/SHANK complex at the PSD (Naisbitt et al., 1999; Tu et al., 1999). The C-terminal SAM domain is a multimerisation domain and is essential for SHANK’s scaffolding function at the PSD (Monteiro and Feng, 2017; Sala et al., 2015).

**Figure 2.**
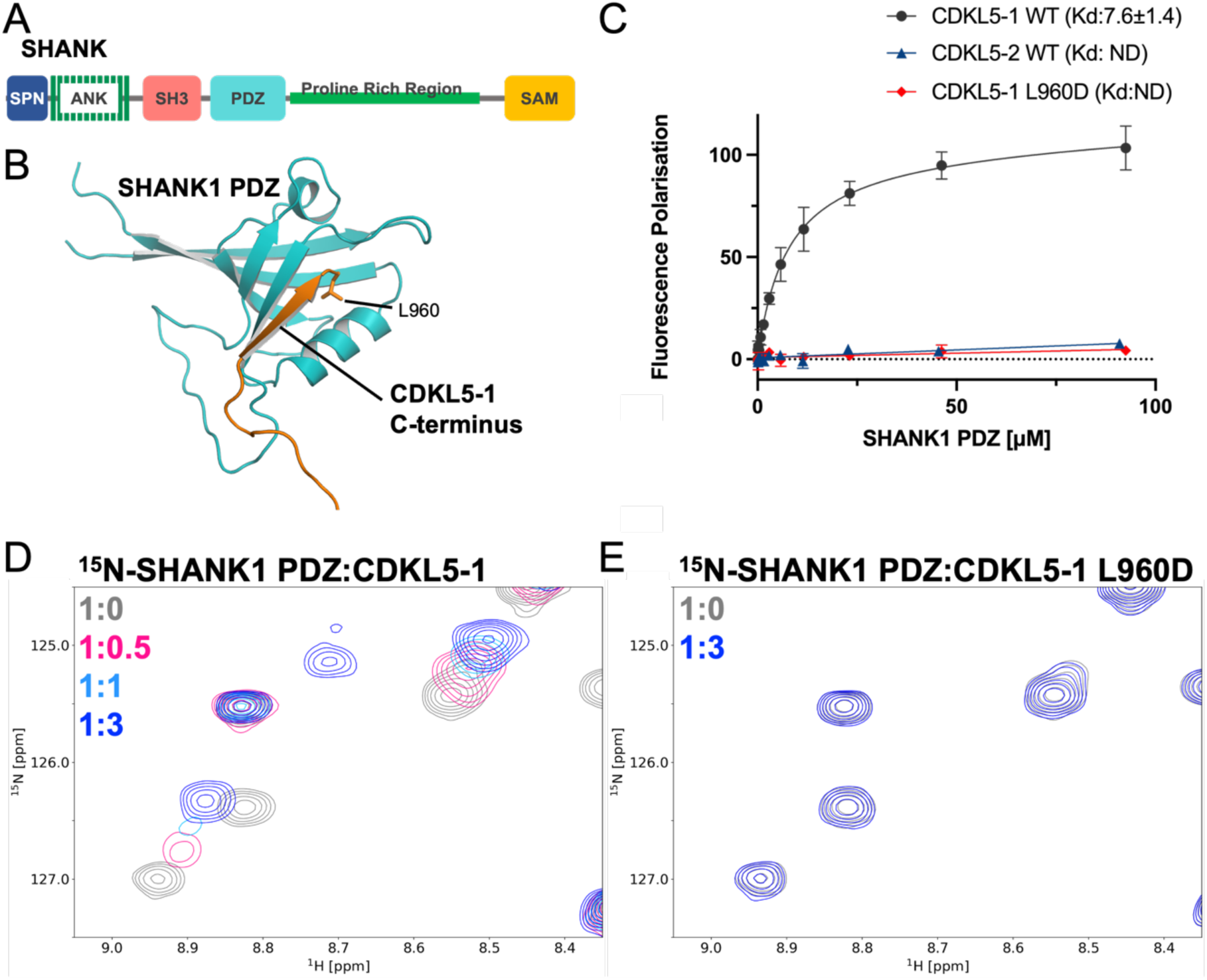
The neuronal isoform of CDKL5 interacts with PDZ domains. **(A)** Domain structure of SHANK proteins, which contain a Shank/ProSAP N-terminal (SPN) domain, an ankyrin repeat region (ANK), an SH3 domain, a PDZ domain, a proline-rich region and a SAM domain. **(B)** AlphaFold predicted structure of SHANK1 PDZ domain (cyan) in complex with CDKL5-1 C-terminal tail peptide (residues 941-960, orange). **(C)** Binding of CDKL5 peptides to the SHANK1 PDZ domain were measured using an FP assay. Fluorescein-labelled CDKL5-1^941-960^, CDKL5-2^1011-1030^ and CDKL5-1^941-960^ L960D peptides were titrated against the SHANK1 PDZ domain. Wildtype CDKL5-1 bound to SHANK1 (black) but CDKL5-2 did not (pink). An L960D mutation of the terminal residue of CDKL5-1 abolished binding (blue). Dissociation constants ± SE (µM) for the interactions are indicated in the legend. All measurements were performed in triplicate. ND, not determined. **(D-E)** NMR titrations of ^15^N-labelled SHANK1 PDZ domain with CDKL5-1 peptides. **(D)** SHANK1 PDZ domain on its own (grey), and with CDKL5-1^941-960^ peptide at 1:0.5 (pink), 1:1 (sky blue) and 1:3 (blue) molar ratios. **(E)** SHANK1 PDZ on its own (grey), and with 1:3 molar ratio of CDKL5-1^941-960^ L960D peptide (blue).

**Figure 3.**
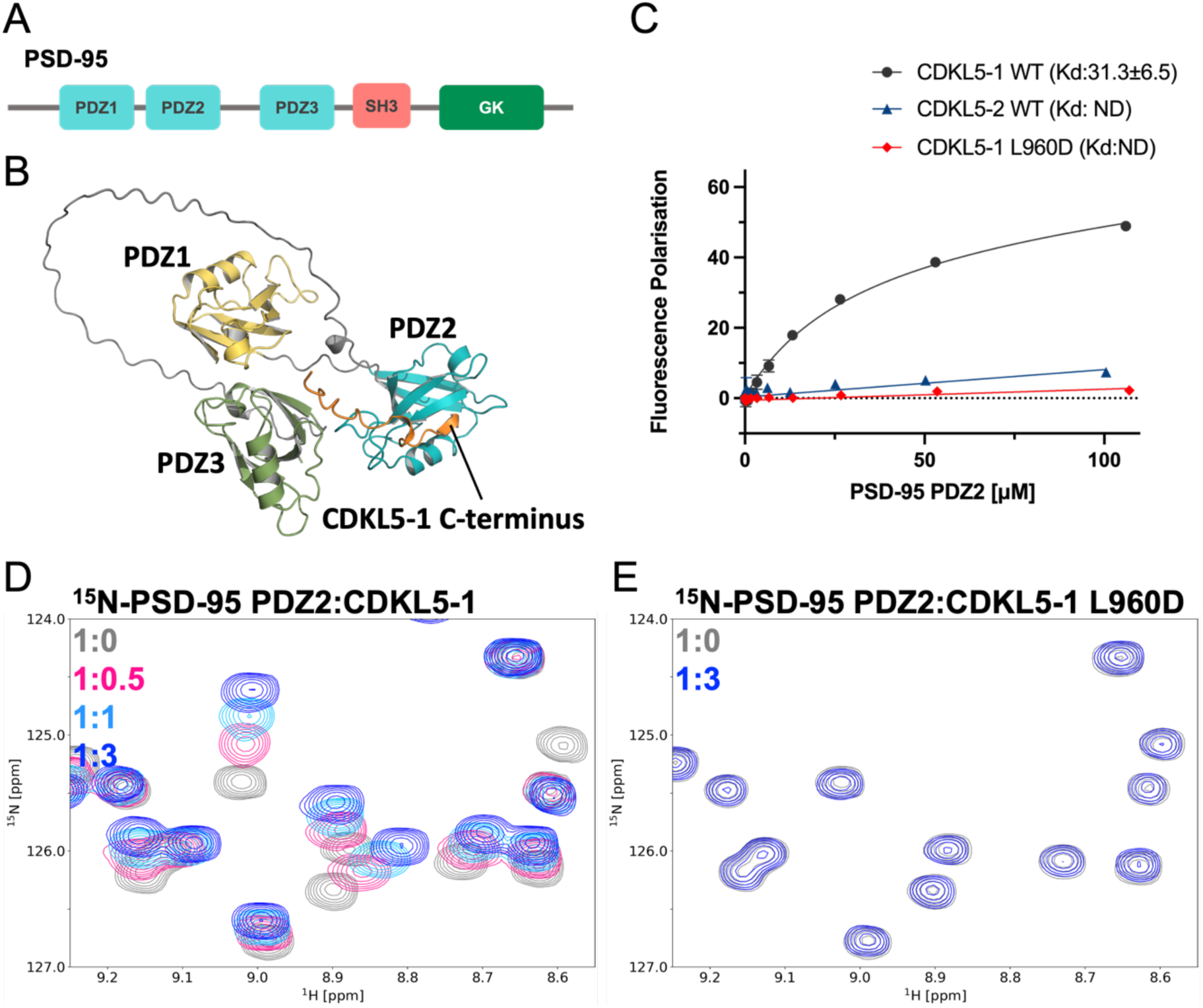
PSD-95 binds the C-terminus of CDKL5 isoform 1. **(A)** Domain structure of PSD-95. PSD-95 has three PDZ domains, an SH3 domain and a guanylate kinase-like (GK) domain. **(B)** AlphaFold predicted structural model of the interaction between PSD-95 PDZ1-PDZ3 (PDZ1; yellow, PDZ2; cyan, PDZ3; green) and the C-terminal tail of CDKL5-1^941-960^ (orange). **(C)** FP assay measuring CDKL5-1^941-960^, CDKL5-2^1011-1030^ and CDKL5-1^941-960^ L960D peptides binding to PSD-95 PDZ2 domain. Dissociation constants ± SE (µM) for the interactions are indicated in the legend. All measurements were performed in triplicate. ND, not determined. **(D-E)** NMR titrations of ^15^N-labelled PSD-95 PDZ2 on its own (grey), and with **(D)** CDKL5-1^941-960^ peptide at 1:0.5 (pink), 1:1 (sky blue) and 1:3 (blue) molar ratios, or **(E)** with CDKL5-1^941-960^ L960D peptide at a 1:3 molar ratio (blue).

CDKL5 localises to the PSD and its colocalisation with PSD-95 and SHANK has been observed in mouse brain (Ricciardi et al., 2012). A connection between CDKL5 and PSD-95 has been detected in multiple proteomics data sets (Murtaza et al., 2022; Unda et al., 2023) and in CDKL5 knockout/knockdown mice, reductions of PSD-95-positive synaptic puncta, decreased spine density and altered spine morphology have all been reported (Della Sala et al., 2016; Lupori et al., 2019; Ricciardi et al., 2012). CDKL5 was previously reported to bind to PSD-95 (Zhu et al., 2013) although the precise molecular basis of this interaction was unclear, appearing to be dependent on PSD-95 palmitoylation. In SHANK3 knockout mouse brains, CDKL5 expression was reduced in striatum (Reim et al., 2017).

### Mechanically active signalling scaffolds in the synapse

Talin is a large mechanosensitive adapter protein that is a component of the synaptic scaffolds that organise each synaptic junction (Ball et al., 2024; Barnett and Goult, 2022; Ellis et al., 2024; Kilinc, 2018). By mediating the adhesive complex that links the actin and microtubule cytoskeletal networks to the integrin family of extracellular matrix (ECM) receptors, talin helps maintain these mechanical couplings. Talin can also mediate cytoskeleton-synapse linkages via direct interaction with the YENPTY motif of Amyloid Precursor Protein (APP) (Ellis et al., 2024). As well as serving as an adapter, talin functions as a mechanically operated signalling scaffold (Ball et al., 2024) due to its 13 helical bundles (rod domains; R1-R13) exhibiting switch-like behaviour. Each talin rod domain can act as a force-dependent binary switch able to reversibly switch between closed (0) and open (1) states depending on mechanical force (Goult et al., 2021, 2013; Yao et al., 2016). The binary switching of each rod domain alters the availability of ligand binding sites and introduces quantised 40-50 nm extensions in the protein length as each rod domain unfolds. As a result, each switching event alters the spatial positioning of subsequent rod domains and proteins interacting with them (Barnett and Goult, 2022). Many proteins engage the talin rod domains (Goult et al., 2021), including Protein Kinase A (PKA) (Kang et al., 2024), and Cyclin-dependent kinase 1 (CDK1) (Gough et al., 2021). The interaction between CDK1 and talin involves a talin-binding leucine-aspartate (LD) motif located in the CDK1 kinase domain which binds directly to talin R8 (Gough et al., 2021). Alignment of kinase domains from other CDKs found that several, including CDKL5, are conserved in this talin-binding region.

In this work, we report that the major neuronal isoform of CDKL5 contains a PDZ-binding motif at its C-terminus that binds to the PDZ domains of SHANKs and PSD-95 tethering CDKL5 to the relatively stable postsynaptic density within the post-synapse. We show that this interaction of CDKL5 with PDZ domains is required for CDKL5 localisation to spines and regulates spine formation. Furthermore, we show that the kinase domain of CDKL5 binds the R8 domain of talin. These two new interactions of CDKL5 provide a new view of how CDKL5 activity may be regulated, whereby mechanical forces on talin lead to changes in the spatial positioning of the kinase domain. Structural transitions in talin R1-R6 domains will move the talin R8-bound CDKL5 kinase domain up and down within the dendritic spine. In this way, we propose a new mechanism of CDKL5 regulation, where the CDKL5 kinase activity is spatially moved up and down within the post-synaptic compartment as a result of neuronal activity controlled mechanical signalling. This leads us to propose a mechanical basis of epilepsy, where misplacement of the CDKL5 kinase domain within the synapse would result in incorrect synaptic signalling, leading to a loss of synchronicity between synapses within the neuronal circuits, resulting in re-entry circuits and erroneous neuronal activity.

## Results

### CDKL5 has a large intrinsically disordered linker region coupling the kinase domain to a highly conserved motif at the C-terminus

The cyclin-dependent kinases comprise a large family of serine/threonine kinases that play central roles in signalling pathways (Kadam et al., 2019; Okuda et al., 2017; Van Bergen et al., 2022). The core CDK family members are relatively small and are comprised of just a kinase domain. However, the CDK-like (CDKL) proteins 1-5 have similar kinase domains but also other regions (Li et al., 2024). CDKL5 has an N-terminal kinase domain (residues 1-306) and a large C-terminal region (residues 307-960 in isoform 1) which AlphaFold (Jumper et al., 2021) predicts is predominantly disordered (Fig. 1A,B).

### CDKL5 has two major isoforms that differ only at their C-terminus

Originally the CDKL5 protein was annotated as a single isoform, residues 1-1030 (UniProt ID:O76039-1), however a second isoform was identified (Williamson et al., 2012), which is now the canonical isoform-1 in UniProt (UniProt ID:O76039-2). Strikingly, this less well studied CDKL5 isoform is the predominant neuronal isoform expressed in brain (Williamson et al., 2012). The two isoforms will hereon in be referred to as CDKL5-1 and CDKL5-2 with CDKL5-1 being the neuronal isoform. The first 905 residues of CDKL5-1 and CDKL5-2 are identical, and this region includes the kinase domain (1-306) and most of the unstructured region (residues 307-905) (Fig. 1B). The C-termini of the two isoforms are different lengths, with CDKL5-2 having 125 residues (residues 906-1030 in CDKL5-2) and CDKL5-1 being shorter, having only 55 residues (residues 906-960 in CDKL5-1).

#### The C-terminus of the neuronal isoform of CDKL5 contains a PDZ domain-binding motif

Conservation analysis using ConSurf (Yariv et al., 2023) revealed that the very C-terminus of CDKL5-1 (residues 951-960) is highly conserved (Fig. 1C). We noticed that this highly conserved region culminates in the residues Glu-Thr-Ala-Leu (-ETAL 957-960) that fits the pattern Ser/Thr-X-Φ-COOH (where X is any amino acid and Φ is any hydrophobic residue) which is the canonical sequence for binding to class I PDZ domains (Songyang et al., 1997). In contrast, the C-terminal sequence of CDKL5-2 culminates in the residues, Leu-Thr-Gly-Lys (-LTGK 1027-1030) which would not be predicted to bind PDZ domains due to the positively charged lysine at the last position.

SHANKs and PSD-95 are major components of the PSD, and both contain PDZ domains, one in each of the three SHANK isoforms (Fig. 2A) and three in PSD-95 (Fig. 3A). We therefore hypothesised that the C-terminal motif in the neuronal isoform of CDKL5 might be binding to PDZ domains to localise CDKL5 activity. If this was the case then the kinase domain would be spatially restrained, anchored via its C-terminus with a zone of activity determined by the unstructured region (Fig. 1D).

We used ColabFold (Jumper et al., 2021) to generate a structural model of a complex between the C-terminal region of CDKL5-1 (residues 941-960) and the PDZ domain of SHANK1 (Fig. 2B) which supported a direct interaction with CDKL5-1^941-960^ bound to the SHANK1 PDZ domain in a canonical fashion. To experimentally test the interaction we performed a Fluorescence Polarisation (FP) assay where fluorescein-labelled CDKL5 C-terminal peptides were titrated with increasing amount of SHANK1 PDZ protein (see Methods and Khan et al., 2021). The FP assay showed a clear interaction between CDKL5-1^941-960^ and SHANK1, with a binding constant (K_d_) of 8.5 µM whereas no interaction was observed with CDKL5-2^1011-1030^ (K_d_ not determinable) (Fig. 2C).

To further explore this interaction, we used Nuclear Magnetic Resonance (NMR) where ^15^N-labelled SHANK1 PDZ domain was titrated with unlabelled CDKL5 peptides. We collected ^15^N-HSQC spectra of ^15^N-labelled SHANK1 PDZ domain on its own and with increasing amounts of CDKL5-1^941-960^ peptide. Addition of CDKL5-1^941-960^ peptide resulted in progressive chemical shifts changes confirming the direct interaction (Fig. 2D).

### A mutation of the terminal residue of CDKL5-1 abolishes binding to PDZ domains

PDZ domains bind to the very C-terminus of their ligands and require that the last residue be hydrophobic to plug into the PDZ domain binding surface (Fig. 2B) (Cabral et al., 1996; Doyle et al., 1996). To test whether the interaction was canonical we designed a CDKL5-1 peptide where the very last amino acid was changed from leucine to aspartic acid, CDKL5-1^941-960^ L960D. The L960D mutant abolished binding as seen by FP assay (Fig. 2C) and NMR (Fig. 2E). The dramatic effect of the C-terminal residue mutation confirms that CDKL5-1^941-960^ is binding canonically to PDZ domains, furthermore it also demonstrates that any modification, or tagging, of the C-terminus of CDKL5 will abolish its interaction with PDZ domains.

### The CDKL5-1 C-terminus binds to the PDZ domains of SHANKs and PSD-95 but binds 5-fold tighter to SHANKs

Having characterised the direct interaction between SHANK1 and CDKL5-1^941-960^ we next wanted to test whether CDKL5-1^941-960^ might also bind the other two SHANK isoforms and PSD-95. Structural modelling of the interaction of the three PDZ domains in PSD-95 with the CDKL5-1^941-960^ peptide robustly positioned CDKL5 bound to both PDZ2 (Fig. 3B) and PDZ3. Using NMR and the FP assay, we tested SHANK2 PDZ, SHANK3 PDZ and PDZ2 and PDZ3 from PSD-95. ^15^N-labelled, SHANK2, PSD-95 PDZ2 and PSD-95 PDZ3 were titrated with CDKL5-1^941-960^ peptide and large chemical shift changes were observed in each PDZ domain spectrum upon addition of peptide (Fig. 3D, Supplementary Fig. 1A and B).

Comparison of the binding affinities of CDKL5-1^941-960^ with SHANKs and PSD-95 showed that the interaction between CDKL5-1^941-960^ and the PDZ domains of SHANKs was ∼5-fold tighter than with the PSD-95 PDZ domains (Fig. 2C, Fig. 3C, Supplementary Fig. 1C-D). The CDKL5-1^941-960^ K_d_ was 7.6 μM for SHANK1 PDZ, 5.2 μM for SHANK2, and 8.6 μM for SHANK3 PDZ compared with 31.3 µM and 40.7 µM for PDZ2 and PDZ3 of PSD-95 respectively (Fig. 2C, Fig. 3C, Supplementary Fig. 1C-D).

No interaction was detectable between CDKL5-2^1011-1030^ and any of the PDZ domains tested (Fig. 2C, 3C, Supplementary Fig. 2) confirming that the C-terminal interaction of CDKL5 with PDZ domains only occurs with isoform 1. Similarly, no interaction was detectable for the CDKL5-1 C-terminal L960D mutant further confirming its efficacy in abolishing binding to PDZ domains (Fig. 3C,E and Supplementary Fig. 3).

### The CDKL5 kinase domain contains a talin-binding site

Talin is anchored to the plasma membrane via the interaction between its head region (FERM) and the cytoplasmic tail of integrins enabling it to function as a mechanosensitive scaffold due to the 13 binary switching rod domains (R1-R13) (Fig. 4A and B). Sequence alignment of the kinase domain region of CDKL5 with other CDKs indicated that the talin-binding site motif we previously identified in CDK1 (Gough et al., 2021) is well conserved in CDKL5, suggesting CDKL5 might also bind to talin R8 domain (Fig. 4C and D, and Supplementary Fig. 4A).

**Figure 4.**
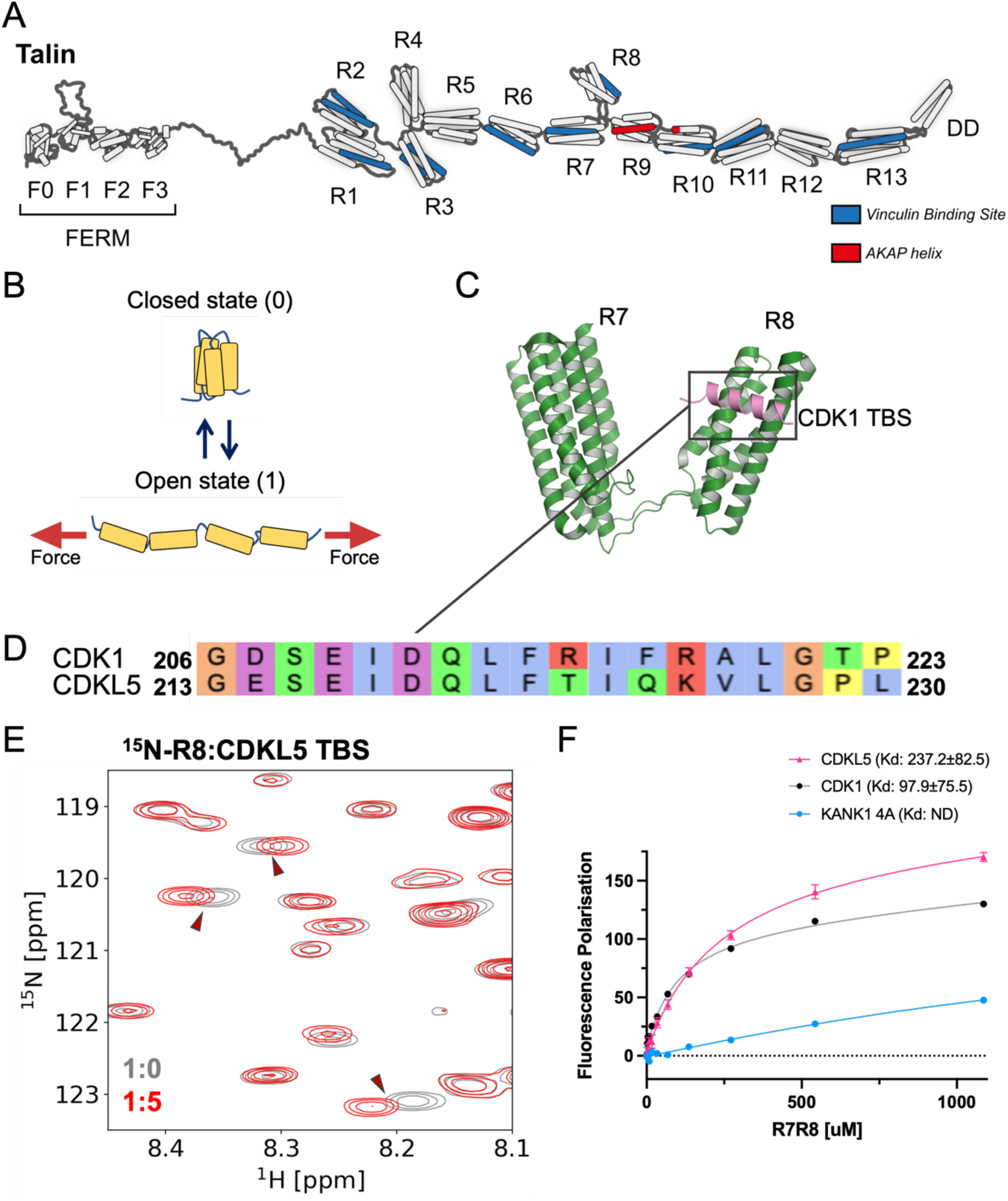
The CDKL5 kinase domain interacts with talin R8. **(A)** Structure model of talin. Talin has an N-terminal head region comprising a FERM domain (F0-F3), 13 rod domains (R1-R13) made of 4- and 5-helix bundles, and a dimerisation domain (DD) at the C-terminus. Helices shown in blue are vinculin binding sites (Wang et al., 2021) and the region showed in red is the PKA binding site (Kang et al., 2024). **(B)** Talin rod domains exhibit switch-like behaviour under force. A helical bundle is shown in the closed (0) and open (1) states. **(C)** The crystal structure of talin1 R7R8 (green) in complex with the CDK1 talin-binding site peptide (pink) (PDB: 6TWN) (Gough et al., 2021). **(D)** Sequence alignment of talin-binding sites from CDK1 (residues 206-223) and CDKL5 (residues 213-230). **(E)** NMR spectra of ^15^N-labelled talin1 R8 on its own (grey), and with CDKL5^213-234^ peptide at a 1:5 ratio (red). **(F)** FP assay measuring talin1 R7R8 binding to the CDKL5^213-234^ peptide (pink) and CDK1^206-223^ peptide (black) against talin1 R7R8. KANK1 4A peptide (blue) was used as a negative control. Dissociation constants ± SE (µM) for the interactions are indicated in the legend. All measurements were performed in triplicate. ND, not determined.

To test this, we collected NMR spectra of ^15^N-labelled talin1 R7R8, R8 and R7 alone and on addition of the putative CDKL5 talin-binding site, CDKL5^213-234^. Chemical shift changes were observed when R7R8 was titrated with CDKL5^213-234^ peptide (Supplementary Fig. 4) which mapped onto the R8 domain (Fig. 4E) and were similar to those seen on addition of CDK1 talin-binding site peptide, CDK1^206-223^ (Gough et al., 2021). In contrast no spectral changes were observed when CDKL5^213-234^ was added to R7 (Supplementary Fig. 4C). We measured the affinity of the CDKL5-talin interaction using the FP assay and found it to be 2.5-fold weaker than CDK1-talin interaction (Fig. 4F).

We next aimed to design a mutation in CDKL5 to disrupt the interaction with talin. In CDK1 talin binding is mediated via hydrophobic residues on one face of the talin-binding site helix (Gough et al., 2021) and these residues are conserved between CDK1 and CDKL5 (Fig. 4D). We mutated two hydrophobic residues in the binding interface, L220 and F221 (L220A, F221A double mutant, 2A) (Supplementary Fig. 5A). This 2A mutant CDKL5 peptide abolished the interaction with talin as shown by NMR (Supplementary Fig. 5B, middle). In addition to the designed 2A mutation, there is a CDD-causing mutation of L220, L220P (Bahi-Buisson et al., 2008; Rosas-Vargas et al., 2008) that we also tested. NMR revealed that L220P significantly weakens the interaction (Supplementary Fig. 5B, bottom).

#### Functional PDZ domain binding of CDKL5-1 is critical for its subcellular localisation

Having identified that the very C-terminus of CDKL5 is essential for interaction with PDZ-domain containing proteins, we next wanted to look at this interaction in cellular contexts. Most studies on CDKL5 have used either the CDKL5 isoform2 or C-terminally GFP- or FLAG-tagged CDKL5 constructs (Bertani et al., 2006; Ricciardi et al., 2012; Rusconi et al., 2008; Dell’Oca et al., 2024; Silvestre et al., 2024), all of which lack a functional PDZ-binding motif. Therefore, we generated a new set of CDKL5 constructs for mammalian cell expression, positioning the GFP-tag at the N-terminus of CDKL5-1; GFP-CDKL5 WT, GFP-CDKL5 L960D and the GFP-tag at the C-terminus, CDKL5-GFP.

We first validated the constructs in cell culture by transfecting the 3 constructs into U2OS cells (Fig. 5). There was a striking difference between the N-terminally and C-terminally tagged constructs, with the N-terminally tagged CDKL5 (GFP-CDKL5 WT) showing punctae throughout the cell which localise proximally to focal adhesions (FAs) (Fig. 5A). In contrast, the L960D mutation (Fig. 5B) and the C-term GFP (CDKL5-GFP) (Fig. 5C) showed no significant accumulation at adhesions (Fig. 5G,H) with the C-terminally tagged construct being diffuse throughout the cytoplasm with nuclear punctae (Fig. 5C). Strikingly, the L960D mutation had significantly higher nuclear localisation than the N-terminally tagged CDKL5 WT (Fig. 5D,E,F,I,J). The reduced FA localisation indicated that the interactions with both membrane-bound PDZ-domain containing proteins and talin were contributing to the localisation at FAs. Together the localisation studies confirm that the C-terminus of CDKL5 is crucial for its localisation.

**Figure 5.**
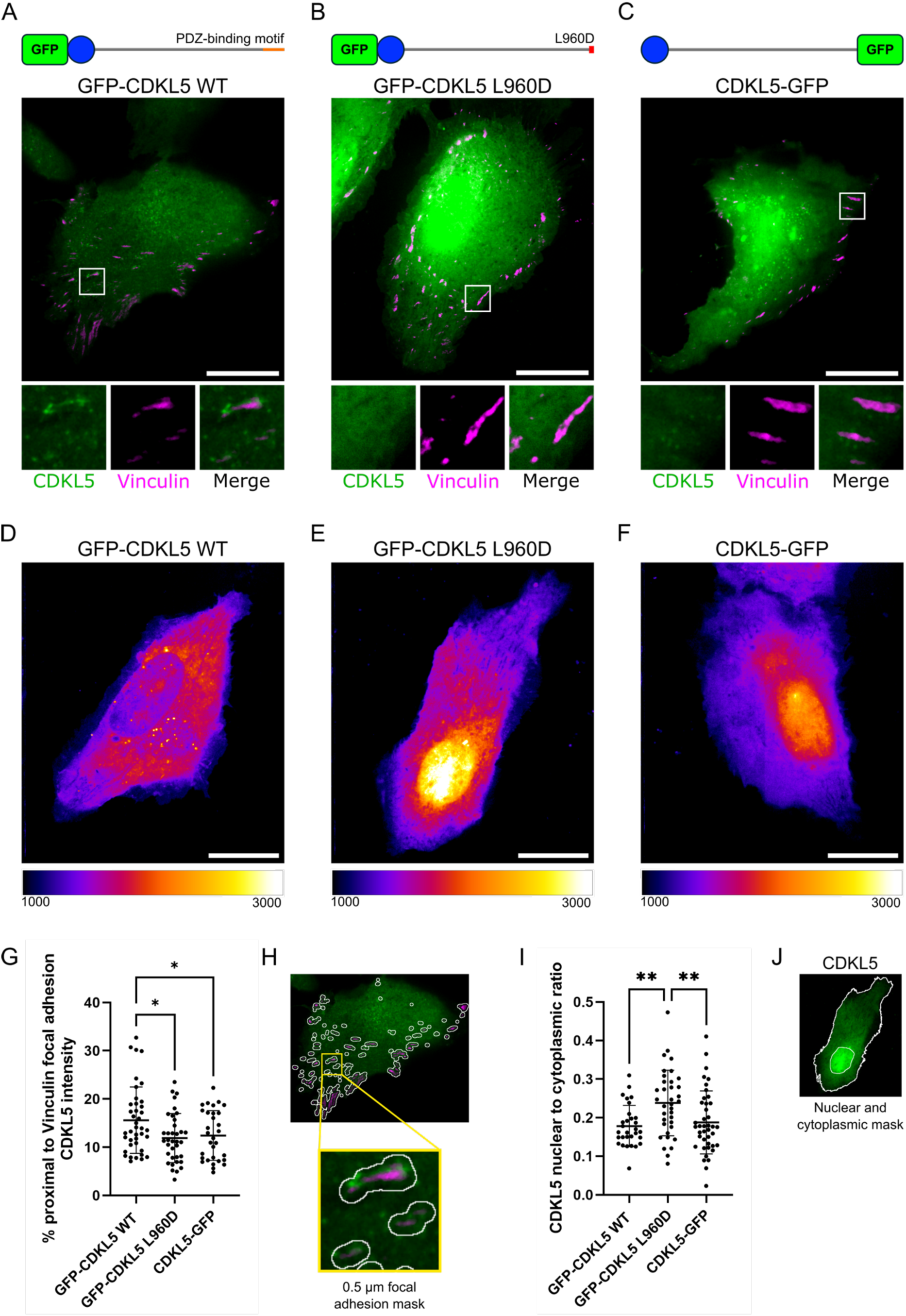
Localisation of CDKL5 to focal adhesions (FAs) requires the PDZ-binding motif. Schematics of the CDKL5 constructs tested are shown above; kinase domain (blue), the PDZ-binding motif (orange) and GFP (green). **(A-C)** Merged images of 2-slice sum projections of U2OS cells co-transfected with mCherry-vinculin (magenta) and **(A)** GFP-CDKL5, **(B)** GFP-CDKL5 L960D or **(C)** CDKL5-GFP (green). Scale bar: 20 μm. Inset panels show the GFP construct localisation around a representative FA. **(D-F)** Merged images of GFP intensity of **(D)** GFP-CDKL5, **(E)** GFP-CDKL5 L960D or **(F)** CDKL5-GFP. **(G-H)** Localisation of the different CDKL5 constructs around FAs. Mean ± SD percentage of GFP construct within a 0.5 μm radius of the mCherry-vinculin FA from a 2-slice sum projection. 31-40 cells, N=5. **(H)** Mask used to quantify GFP puncta around FA. **(I-J)** Nuclear localisation of the different CDKL5 constructs. Mean ± SD nuclear to cytoplasmic ratio of a sum projection of each GFP construct 30-40 cells, N=5. **(J)** Mask used to quantify GFP nuclear to cytoplasmic ratio. Significance test used was an ordinary one-way ANOVA with Sidak’s multiple comparisons tests: * p ≤ 0.05, ** p ≤ 0.01.

#### The PDZ-binding motif anchors CDKL5 to the postsynaptic density in primary neurons

We next wanted to assess whether the PDZ-binding motif is important for CDKL5 localisation in neurons. For this, we transfected rat primary cortical neurons with CDKL5 constructs and imaged the neurons using confocal microscopy. Similar to U2OS cells, the N-terminally tagged CDKL5 (GFP-CDKL5 WT) exhibited a striking difference in localisation compared to the L960D mutant with disrupted PDZ-binding motif (GFP-CDKL5 L960D) and the C-terminally tagged CDKL5 (CDKL5-GFP). The N-terminally tagged CDKL5 accumulated in focalised puncta along the dendritic shaft and in spines, whereas the GFP-CDKL5 L960D and the C-terminally tagged CDKL5 exhibited more diffuse localisation (Fig. 6A). This indicates that a functional PDZ-binding motif is important for CDKL5 localisation along the dendrites.

**Figure 6.**
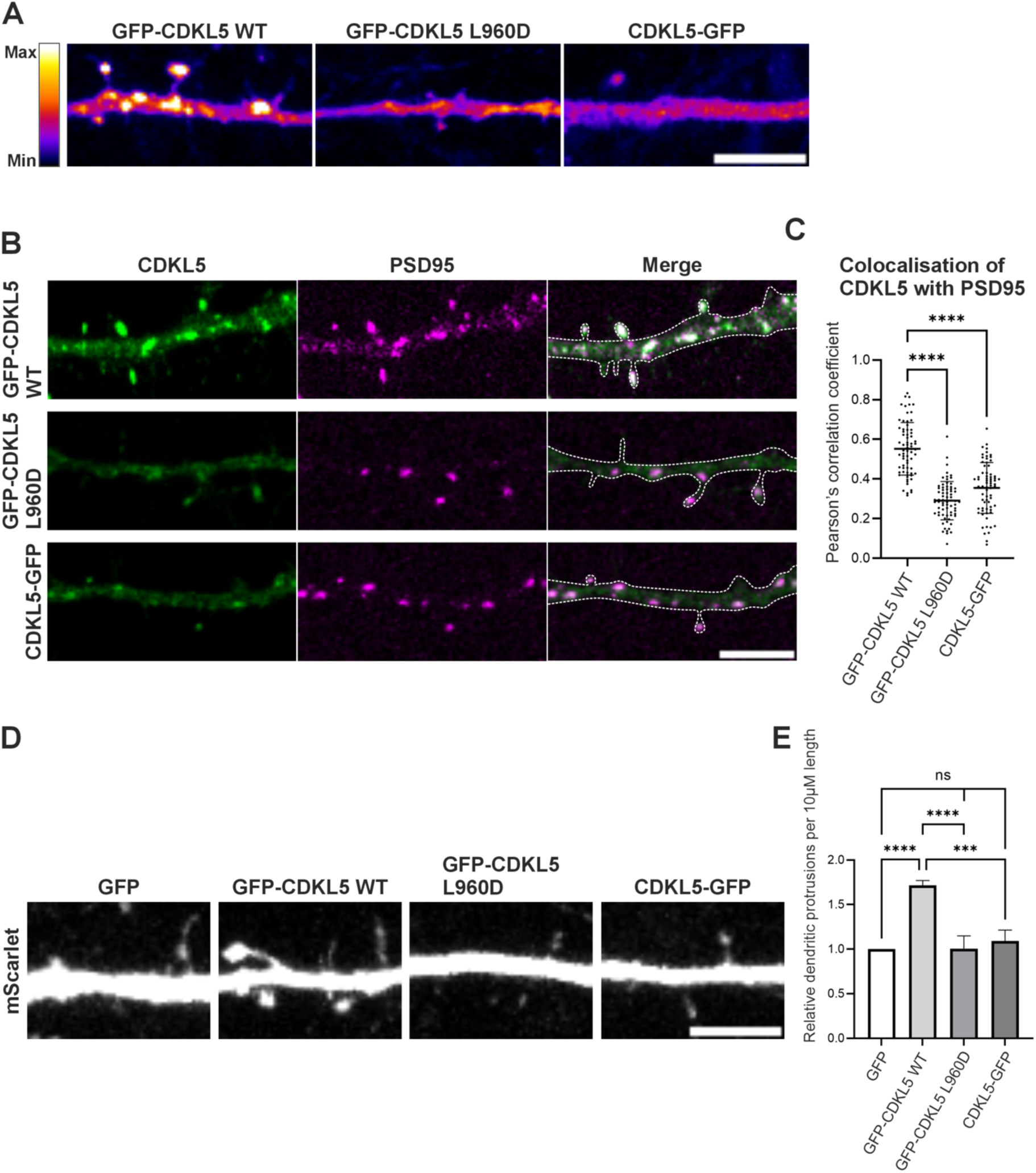
A functional PDZ-binding motif is important for the localisation and activity of CDKL5 at postsynaptic sites. DIV15 primary cortical neurons were transfected with; **(A)** GFP-CDKL5 WT (first panel), GFP-CDKL5 L960D (second panel), and CDKL5-GFP (third panel). The GFP signal is shown in Flame pseudo-colour LUT. Left: Flame LUT maximum and minimum. **(B)** CDKL5 constructs and PSD-95-pTagRFP. GFP-CDKL5 WT (first row), GFP-CDKL5 L960D (second row), and CDKL5-GFP (third row). CDKL5 constructs are in green (first column), PSD-95 is in magenta (second column), and the merged signal represents the degree of colocalisation (third column). The white dotted line indicates the boundary along the dendritic shaft. **(C)** The extent of CDKL5 colocalisation with PSD-95 in the dendrites was quantified and represented as Pearson’s correlation coefficient. **(D)** CDKL5 constructs and mScarlet. GFP-transfected neurons served as controls (first panel), GFP-CDKL5 (second panel), GFP-CDKL5 L960D (third panel), and CDKL5-GFP (fourth panel). A section of the dendritic shaft is shown. The mScarlet signal is shown in greyscale. **(E)** The relative number of dendritic protrusions per 10 µm of dendritic shaft is represented as a histogram. (C and E) Statistical analysis between the groups was performed using ordinary one-way ANOVA with multiple comparisons: **** p ≤ 0.0001, *** p < 0.001, ns – not significant. Scale bars: 5 µm

Given that CDKL5 interacts with PSD-95 through its PDZ-binding motif, we tested whether this motif targets CDKL5 to the postsynaptic density. Primary cortical neurons were co-transfected with PSD-95-pTagRFP and either GFP-CDKL5, GFP-CDKL5 L960D, or CDKL5-GFP. The N-terminally tagged CDKL5 (GFP-CDKL5) strongly colocalised with PSD-95 along the dendritic shaft and spines (Fig. 6B, first row). In contrast, the PDZ-binding mutant version of CDKL5 (GFP-CDKL5 L960D, Fig. 6B, second row) and the C-terminally tagged CDKL5 (CDKL5-GFP, Fig. 6B, third row) showed significant reduction in colocalisation, as quantified by Pearson’s correlation coefficient (Fig. 6C). These findings underscore the importance of the CDKL5 PDZ-binding motif and its functional interaction with PSD-95 for the correct targeting of CDKL5 activity at neuronal synapses.

CDKL5 knockdown has been reported to result in a reduced number of spines in primary hippocampal neurons (Mottolese et al., 2025; Ricciardi et al., 2012). Therefore, we wanted to determine whether ectopic CDKL5 expression would promote spine formation by quantifying the density of dendritic protrusions in CDKL5-transfected neurons. To achieve this, CDKL5 constructs were co-transfected into primary cortical neurons along with a morphological marker, mScarlet, which was used to quantify the number of dendritic protrusions (Fig. 6D). Indeed, compared to neurons transfected with an empty GFP vector, the N-terminally tagged GFP-CDKL5 displayed approximately a 70% increase in the number of dendritic protrusions (Fig. 6E). In contrast, the PDZ-binding mutant (GFP-CDKL5 L960D) and CDKL5-GFP transfected neurons showed no significant increase in the number of dendritic protrusions compared to GFP-transfected control neurons. This indicates that the PDZ-binding motif is important for the correct targeting and neuronal activity of CDKL5, mirroring the effects previously observed with CDKL5 downregulation (Mottolese et al., 2025; Ricciardi et al., 2012).

## Discussion

CDKL5 deficiency disorder (CDD) is a devastating condition caused by mutation in the *CDKL5* gene and results in childhood epilepsy with seizures occurring soon after birth, and throughout the child’s life. Development is also impaired in children with CDD. The *CDKL5* gene is located on the X chromosome and produces the kinase, CDKL5 which is essential for normal brain development and synaptic activity.

Here we report two new protein-protein interactions of CDKL5 which identify a new spatial model of regulation of CDKL5 activity. We identify i) a PDZ-binding motif at the very C-terminus of the neuronal isoform of CDKL5 that interacts with SHANK proteins and PSD-95 and ii) a talin-binding motif in the CDKL5 kinase domain which interacts with talin R8. We show that the interaction with PDZ domains is required for correct recruitment of CDKL5 to the postsynaptic density and that disruption of the PDZ-binding motif in CDKL5 results in its mislocalisation and impaired spine formation. As the C-terminus of CDKL5 is anchored, we propose a novel role for the large unstructured region in spatially restricting CDKL5 activity, by acting as a tether for the CDKL5 enzymatic domain (Fig. 7A). Targeting of CDKL5 to the synapse through the PDZ-binding motif at the C-terminus, tethering of the kinase domain via the unstructured region, and positioning of the kinase domain through interaction with talin R8 leads to a new view of CDKL5 as a mechanically-regulated kinase (Fig. 7B).

**Figure 7.**
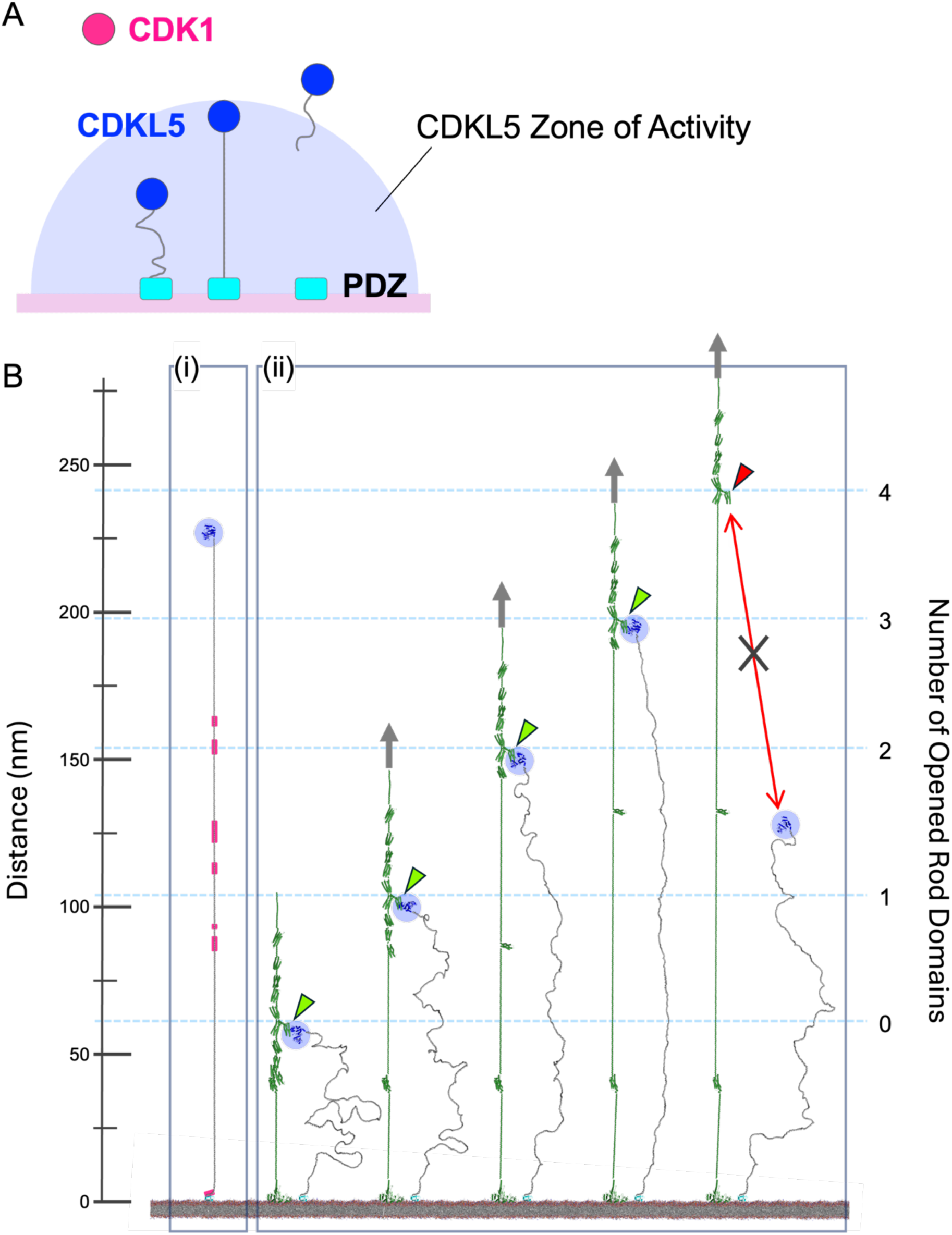
Conceptual model of CDKL5 as a mechanically-operated kinase. **(A)** The PDZ-binding motif tethers the CDKL5 at the synapse, and the large unstructured region defines a “zone of activity” in which the kinase domain is active (blue semi-circle), with radius the length of the tether (>200 nm). Epilepsy-causing mutations that introduce premature stop codons untether CDKL5. CDK1 is not tethered. **(B)** (i) CDKL5 is shown fully extended with the C-terminus bound to membrane-tethered PDZ domains (cyan). The kinase domain is highlighted via a blue sphere, and highly conserved regions on the CDKL5 unstructured region are highlighted (magenta). The scale bar on the left is the distance from the membrane in nanometres. (ii) The CDKL5 kinase domain can be moved up and down in the synapse by talin domain switching (green). Talin is shown membrane-tethered via its interaction with integrins and/or APP. The distance of R8 from the membrane varies depending on the number of rod domains from R1-R6 that are unfolded. The scale bar on the right and the dashed lines indicate the position of R8 from the membrane with (left to right): all rod domains folded; one rod domain (R3) unfolded; two rod domains (R3 and R5) unfolded; three rod domains (R2, R3, and R5) unfolded; and four rod domains (R2, R3, R5, and R6) unfolded. The kinase domain of CDKL5 (blue) binds to R8 only when folded R8 exists within the range the kinase domain can reach which is governed by the length of the unstructured region of CDKL5. Green arrows: R8 bound to CDKL5 kinase domain; red arrow: R8 not interacting with CDKL5 kinase domain. The connections to the force generating machinery are shown by grey arrows which point in the direction of the force vector. See also [Supplementary Movie 1](https://www.youtube.com/watch?v=WC6t-A5GHPo)

### Discovery of a canonical interaction between CDKL5 and PDZ domains

We report how the CDKL5 PDZ-binding motif can bind canonically to the PDZ domains of PSD-95 and SHANK proteins. Disruption of this interaction leads to increased CDKL5 nuclear localisation in cells (Fig. 5) and loss of CDKL5 synaptic localisation in neurons (Fig. 6). An interaction between CDKL5 and PSD-95 was reported previously (Zhu et al., 2013), but the biochemical elucidation of the interaction was complicated by the fact the isoform used lacked the PDZ-binding motif. There are two CDKL5 isoforms, that have different expression patterns (Williamson et al., 2012), and the two isoforms differ only at the C-terminus, with only isoform 1, the major neural isoform, containing a PDZ-binding motif (sequence -ETAL) (Fig. 1B). This PDZ-binding motif is clearly evident in most mammalian *CDKL5* genes including human (UniProt accession number: O76039), mouse (UniProt accession number: A0A0G2JGW6) and rabbit (UniProt accession number: A0A5F9CN53). However, the annotation of the rat *CDKL5* gene still lacks the PDZ-binding motif despite the PDZ-binding motif having been confirmed in rat (Williamson et al., 2012), and being listed on NCBI (Accession number: XP_002727631.1). It is possible that the lack of the PDZ-binding motif in rat CDKL5 may be an annotation issue as all other mammals we checked contain a CDKL5 isoform with this sequence.

### Previous neuronal studies on CDKL5 where the C-terminus has been altered will have looked at mislocalised protein

Many studies have investigated CDKL5 localisation and function but have used either i) the CDKL5 isoform which lacks the PDZ-binding motif (Bertani et al., 2006; Ricciardi et al., 2012; Rusconi et al., 2008) or ii) a C-terminal tag (Dell’Oca et al., 2024; Silvestre et al., 2024), presumably tagging the C-terminus to position the tag away from the N-terminal kinase domain. However, our work identifies that these studies will have inadvertently abolished the PDZ-binding motif with the result that CDKL5 will be mislocalised. Isoform 1 is also the predominantly expressed CDKL5 isoform in other human tissues (Williamson et al., 2012) so non-neuronal studies on CDKL5 will also have been impacted if the wrong isoform or tagging strategy was used.

Overexpression of CDKL2 was recently shown to phosphorylate the CDKL5 substrate, EB2 in CDKL5 knockout mouse neurons leading to the suggestion that CDKL2 overexpression might represent a potential therapeutic strategy for CDD (Silvestre et al., 2024). However, C-terminally FLAG-tagged CDKL5 and CDKL2 were used to compare EB2 phosphorylation levels, and the C-terminal tagging of CDKL5 will have detached CDKL5 from its proper localisation. Furthermore, CDKL2 does not have a PDZ-binding motif so is unlikely to be spatially regulated at the synapse in the way CDKL5 is. Additionally, the talin-binding site in CDKL5 is poorly conserved in CDKL2 and so is unlikely to bind to talin. It is possible that CDKL2 could compensate for loss of function of CDKL5 isoform 2 which lacks the PDZ-binding motif but less likely for loss of function of isoform 1, the major isoform in the brain.

### A proposed model of CDKL5 as a mechanically-operated kinase

The discovery of an interaction between the CDKL5 kinase domain and the R8 domain of talin indicates that the localisation of the kinase domain and CDKL5 activity could be spatially positioned by the mechanical unfolding/refolding of preceding talin rod domains as a function of neuronal activity. Our previous work on the interactions between talin and Protein Kinase A (PKA) (Kang et al., 2024) and CDK1 (Gough et al., 2021) provide two paradigms for how kinases can be mechanically operated through talin. PKA binds to talin only when the 9^th^ switch, R9 is in the unfolded (1) state, upon which the AKAP (A-kinase anchoring protein) helix is exposed and can scaffold cAMP-signalling at a precise location along the talin scaffold (Kang et al., 2024). On the other hand, CDK1 binds to talin only when the 8^th^ switch, R8 is in the folded (0) state (Gough et al., 2021). In both cases the enzymatic scaffolding is dependent on the talin switch patterns which are the result of mechanical signals acting on talin.

CDKL5 provides a new paradigm for how mechanically-operated kinase might work in that it binds to talin R8 in a similar way to how CDK1 binds (Gough et al., 2021). However, a novel feature of CDKL5 is the tethering of the kinase domain to the membrane which identifies two specific roles for the unstructured region as i) a tether, preventing CDKL5 diffusing away from the synapse and ii) dictating the zone in which CDKL5 activity will be retained. In this way, the PDZ-binding motif would anchor CDKL5 at the synapse and the interaction of CDKL5 with talin R8 would permit precise localisation of the CDKL5 kinase activity (Fig. 7)

### Re-evaluation of CDD-causing CDKL5 mutations in the mechanically-operated kinase model of epilepsy

Many of the identified CDKL5 mutations that cause CDD introduce premature stop codons into the protein (Fig. 8A). We show how even a single point mutation, L960D of the last residue abolishes the interaction with PDZ domains and leads to loss of localisation of CDKL5 activity at synaptic sites (Fig. 6) and increased nuclear localisation (Fig. 5). Premature stop codons would detach CDKL5 from the anchoring at the PSD and lead to mis-localised kinase activity (Fig. 8). The unstructured C-terminal region has been reported to regulate CDKL5 localisation and activities (Lin et al., 2005; Rusconi et al., 2008) and CDKL5 nuclear/cytosol localisation is precisely altered over neuronal developmental stages in mice brain suggesting that CDKL5 functions specifically at each site (Rusconi et al., 2008). It is possible that, during development, levels of the two isoforms are tightly regulated to enable correct levels of CDKL5 in the nucleus and at the forming synapses. Premature stop codon mutants would lose the targeting sequence from isoform 1, making it similar to isoform 2, and disrupting the precise regulation of CDKL5 localisation and activity required for neuronal development.

**Figure 8.**
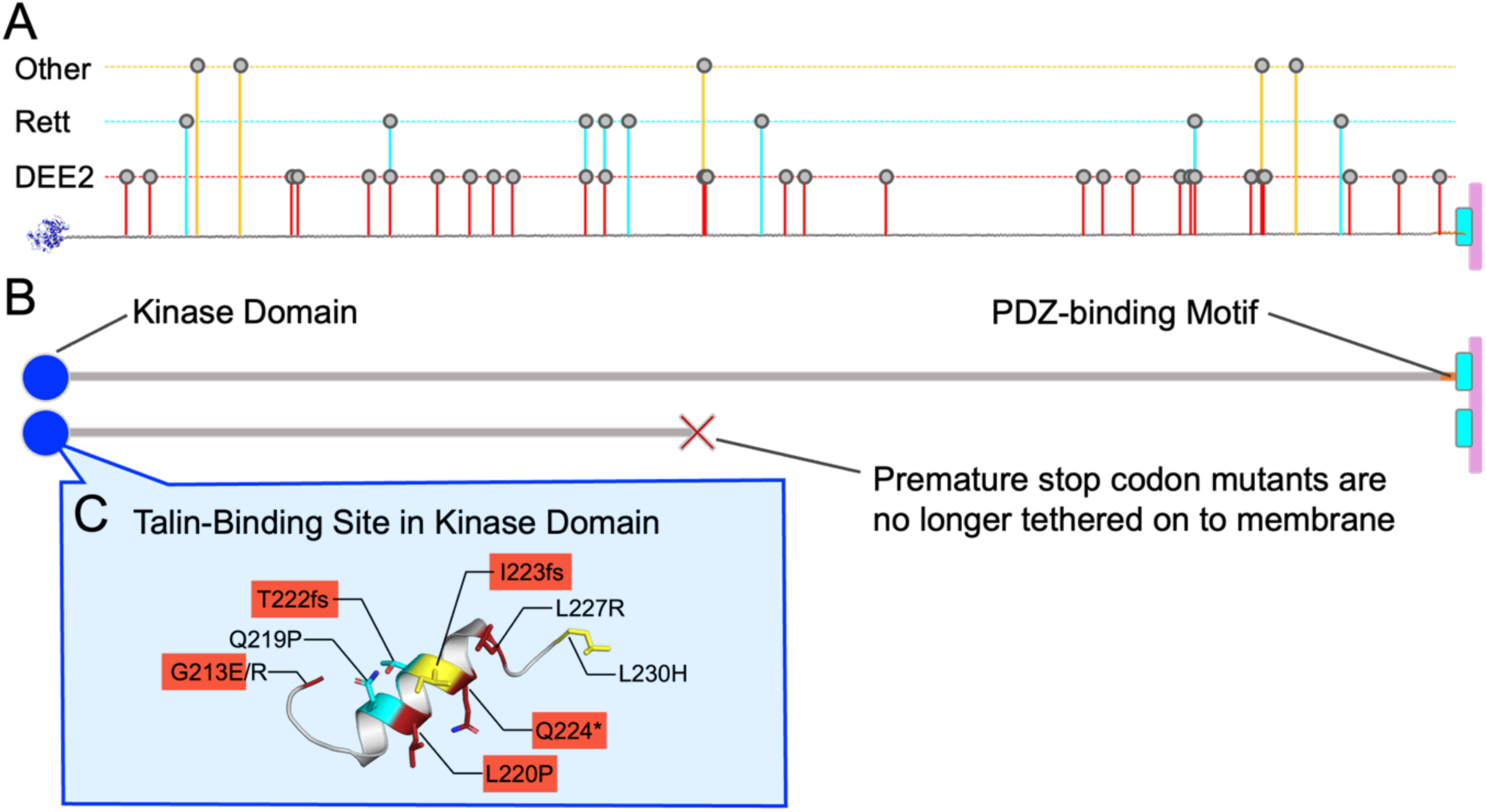
Pathogenic CDKL5 missense variants in the unstructured region uncouple CDKL5 from the synaptic membrane. **(A)** The unfurled structural model of CDKL5 is shown. The kinase domain (blue) is shown to the left, and the PDZ-binding motif (red) is shown engaging PDZ domains (cyan) at the synaptic membrane (pink). Lollipops indicating the stop codon mutations reported as pathogenic (red; DEE2, cyan; Rett, and yellow; other) in the CDKL5 unstructured region are shown. **(B)** Model of CDKL5 full length (top) and with a stop codon mutation in the middle of the unstructured tail (bottom). Any premature truncation will have the same consequence that the kinase domain (blue) is no longer tethered onto the plasma membrane. **(C)** There are many CDKL5 missense variants in the kinase catalytic domain that effect its activity. A subset of these variants map onto the talin-binding site (red; DEE2, cyan; atypical Rett, and yellow; other disease). fs indicates frame shift mutation, * indicates stop codon mutation. Data taken from ClinVar (Landrum et al., 2014) and accession numbers used are given in the Methods.

There are also many CDD-causing mis-sense mutants in CDKL5 which cluster in the kinase domain, and likely impact on kinase activity itself. We note that there are a number of mis-sense mutations in the talin-binding site (Fig. 8C) and L220P impacts on the interaction with talin (Supplementary Fig. 5). Mis-sense mutations in the unstructured region might be less deleterious as in our model the role of this region is as the tether connecting the kinase to the membrane. However, premature stop codons, irrespective of where on the unstructured region, would be deleterious as they would uncouple CDKL5 from its anchor points leading to its mislocalisation and accumulation in the nucleus.

### CDKL5 as a mechanically regulated signalling scaffold

We provide a model for how the CDKL5 kinase domain is positioned in the synapse by its interaction with the mechanosensitive signalling scaffold talin (Fig. 7). However, what is striking is that as CDKL5 is anchored, movement of the kinase domain up and down in the synaptic bouton will lead to the extension of the CDKL5 unstructured region (Fig. 7). Analysis of the conservation of CDKL5 through evolutionary time using ConSurf (Yariv et al., 2023) identifies multiple ultra conserved regions of unknown function in the unstructured region (Fig. 7B, and Supplementary Fig. 6). These regions almost certainly represent binding motifs for other, as yet unidentified, proteins. Therefore, as the linker region gets stretched out it will likely be serving to spatially position these interacting proteins in the z-dimension. As a result, our data indicates a putative new role of CDKL5 as a mechanically regulated signalling scaffold itself. Defining which neuronal proteins engage with these binding-motifs will be a crucial next step in a better understanding of CDKL5 function.

### Future directions

We define here how CDKL5 is anchored at the postsynaptic site. Future work should look at whether CDKL5 is also being anchored at the presynaptic site, and if so to what protein(s). There are many PDZ domains in the brain, and members of the DLG (disks large)-MAGUK subfamily, to which PSD-95 belongs, such as SAP-97 (synapse-associated protein 97, also known as DLG1; disk large homolog 1) have class I PDZ domains and are found at both presynaptic and postsynaptic sites (Aoki et al., 2001).

Furthermore, our work identifies CDKL5 itself as a mechanically-operated signalling scaffold. As the unstructured region is stretched out proteins bound to these sites will get organised in the z-dimension away from the PSD (Fig. 7B). Our conservation analysis identifies a number of highly conserved sequences in the unstructured region of CDKL5 that likely define binding sites for currently unidentified ligands (Fig. 7, Supplementary Figure 6). Therefore, a detailed proteomic analysis of this region of CDKL5 will be a high priority to identify new components of CDKL5 biology and pathophysiology.

The link between CDKL5, an established synaptic enzyme involved in healthy brain functioning, and the mechanosensitive signalling scaffold talin provides more evidence in support of the MeshCODE theory of a mechanical basis of memory (Barnett and Goult, 2022; Goult et al., 2021) centred on the idea that the binary switches in talin might allow persistent patterns of information to be written into the shape of the talin molecules (Ball et al., 2024; Goult, 2021; Goult et al., 2021; Yao et al., 2016). As these switch patterns control signalling outputs, they represent a potential way to encode information in the form of memories in a binary format (Barnett and Goult, 2022; Goult et al., 2021). Our recent discovery of a direct link between talin and Amyloid Precursor Protein (APP) which links talin to memory loss and Alzheimer’s Disease (Ellis et al., 2024) further supports this concept. Future work should look at the role of talin in epilepsy and in visualising the changing locations of talin-bound enzymes within the synapse.

In summary, we define two new interactions of CDKL5 that provide a new view of how it is regulated, and importantly how it is misregulated in epilepsy. We hope that this information will lead to a deeper understanding of CDKL5 and talin function within dendritic spines. We predict that the CDKL5 kinase domain will move up and down within the synapse in response to mechanical folding/unfolding events in talin. The precise positioning of CDKL5 within synapses will be key to its function, and mislocalisation at one synapse will lead to the desynchronisation of that synapse relative to other synapses in the neuronal circuit. We envisage that this desynchronisation of one synapse relative to another due to out of context enzymatic activity will lead to re-entry circuits and epilepsy.

## Methods

### Protein expression and purification

Recombinant talin, PSD-95 and SHANK proteins were expressed from pET151 plasmids containing; mouse talin1 (UniProt: P26039) R7R8 (residues 1357-1659), R7 (residues 1357-1653 Δ1454-1586) and R8 (residues 1461-1580); mouse talin2 (UniProt: B2RY15) R7R8 (residues 1360-1656), and codon optimised synthetic genes (GeneArt; Life Technologies) for human PSD-95 (UniProt: P78352) PDZ2 (residues 155-249), and PDZ3 (residues 302-402), mouse SHANK1 PDZ (UniProt: D3YZU1, residues 654-762), mouse SHANK2 PDZ (UniProt: Q80Z38, residues 247-340), and mouse SHANK3 PDZ (UniProt: Q4ACU6, residues 570-664).

BL21(DE3) Star *E. coli* cells were transformed with the individual plasmids. As starter cultures, these cells were grown in 10 mL of LB + 100 µg/mL ampicillin at 37°C overnight for inoculation into 750 mL scale culture. To produce ^15^N-labelled protein, 2M9 minimal medium containing 1 g/L of ^15^N-ammonium chloride, 4 g/L of glucose and 100 µg/mL ampicillin was used. Once an OD_600_ of 0.7-0.8 was reached the cultures were induced with 1 mM IPTG and incubated at 20°C overnight. Cells were harvested by centrifugation at 6000 RCF for 10 minutes and resuspended in 20 mM Tris-HCl pH 8, 500 mM NaCl and 20 mM imidazole. Nickel affinity chromatography was performed using a 5 mL HisTrap HP column (Cytiva). Proteins were dialysed into 20 mM Tris-HCl pH 8, 50 mM NaCl with AcTEV protease (Invitrogen) to remove the His-tag. A 5 mL HiTrap Q HP anion exchange column (Cytiva) was used for further purification. More details on protein preparation are described in (Khan et al., 2021).

### Fluorescence Polarisation Assay (FP)

The following peptides were synthesised by GLBiochem (Shanghai)

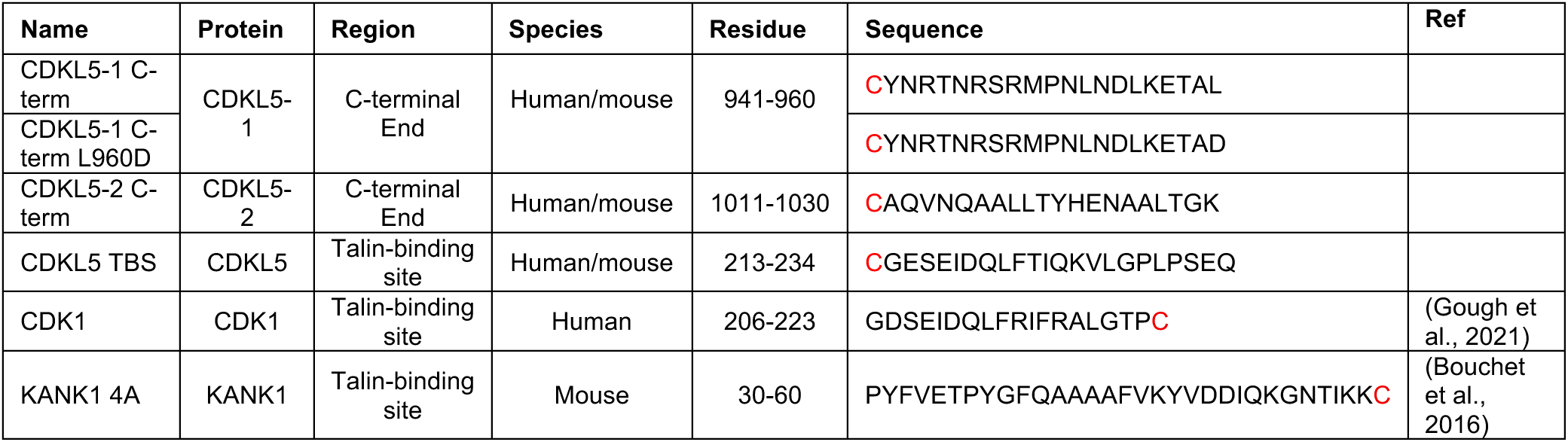

Synthetic peptides were designed with a single non-native N-terminal cysteine to couple with maleimide-fluorescein dye (Thermo Fisher Scientific). Target proteins were prepared with a twofold serial dilution in PBS (10 mM Na_2_HPO_4_, 1.8 mM KH_2_PO_4_, pH7.4, 137 mM NaCl, 2.7 mM KCl). CLARIOstar (BMG LabTech) or HIDEX Sense Microplate reader (HIDEX) were used to measure the polarisation at 25°C (excitation: 482 ± 8 nm; emission: 530 ± 20 nm, or 485 ± 10 nm; emission: 535 ± 20 nm for fluorescein and excitation: 544 ± 20 nm; emission: 590 ± 20 nm for Bodipy). Data were fitted to a one-site total binding equation in GraphPad Prism 8.

### Nuclear Magnetic Resonance (NMR)

Protein samples for NMR were prepared in 12 mM NaH_2_PO_4_, 6 mM Na_2_HPO_4_, pH 6.5, 50 mM NaCl, 2 mM DTT, 5% (v/v) D_2_O at 150 µM final concentration and titrated with peptides in identical buffer. A Bruker Avance III 600/700 MHz NMR spectrometer equipped with CryoProbe was used to collect NMR spectra at 298K. The data were processed and analysed using TopSpin and CCPN Analysis (Skinner et al., 2015).

### Cloning

The template for CDKL5-2 was sourced from p663-UBC-miniTurbo-V5-CDKL5_IDG-K which was a gift from Ben Major (Addgene plasmid #170578; http://n2t.net/addgene:170578; RRID: Addgene_170578). The template for CDKL5-isoform1 (residues 746-959) was ordered as a synthetic gene (GeneArt; Life Technologies Ltd).

GFP-CDKL5-2 (1-1030)-WT was constructed by subcloning the CDKL5-2 sequence into the XhoI/BamHI sites in eGFP-C1. The CDKL5-1 constructs were prepared by initially subcloning residues 1-745 of isoform 2 into the XhoI/KpnI sites of either eGFP-C1 or eGFP-N1. A second round of subcloning of the isoform 1-specific C-terminus into the KpnI/BamHI sites of the previous plasmids was performed. The C-terminal residue of CDKL5-1 was specified in the reverse primer (L960 or the mutated D960). The GFP-CDKL5 kinase domain construct (residues 1-311) was prepared by subcloning into the XhoI/BamHI sites of eGFP-C1.

Whole plasmid sequencing was performed by Plasmidsaurus using Oxford Nanopore Technology with custom analysis and annotation to verify all constructs. The five constructs generated for this study have been deposited in Addgene (https://www.addgene.org/Ben_Goult/).

### Cell culture

Human osteosarcoma U2OS cells (HTB-96™) were obtained from ATCC®. They were maintained in GlutaMAX™ DMEM (Gibco, Thermo Scientific, 31966047) with 10% fetal bovine serum (Sigma-Aldrich) at 37°C and 5% CO*_2_* and were used for a maximum of 20 passages.

### DNA transfection into U2OS cells

Cells were seeded at ∼4.5 x 10^5^ cells per well in a 35 mm glass bottom dishes (CellVis, D35-14-1.5N) pre-coated with 50 μg/mL collagen and 10 μg/mL fibronectin (Sigma-Aldrich, FL1141) between 16–24 hours prior to transfection. Cells were co-transfected with mCherry Vinculin and either GFP-CDKL5, GFP-CDKL5 L960D or CDKL5-GFP 24 hours before imaging using Lipofectamine 3000 and ∼2 μg of each plasmid for all transfections (∼4 μg total).

### Primary cortical neuron culture and transfections

Cortical neuronal cultures were prepared as described previously (Sahu et al., 2019). Briefly, cortices were dissected from Wistar E17-E18 rat embryos. The tissues were enzymatically dissociated to obtain neurons, which were plated on glass coverslips coated with 0.1 mg/mL poly-L-lysine (PLL) (Sigma-Aldrich) and cultured in MACS® Neuro Medium (Miltenyi Biotec) supplemented with MACS® NeuroBrew®-21 (Miltenyi Biotec), L-glutamine (Invitrogen), and penicillin-streptomycin (Gibco). Every 7 days, half of the cell culture medium was replaced with fresh MACS medium with supplements. All neurons were transfected on DIV14 and fixed at DIV15.

At DIV14, 100,000 cortical neurons per coverslip were transfected with 2 µL of Lipofectamine 2000 (ThermoFisher) and 1.2 µg of DNA following the manufacturer’s recommendations. The plasmids used for transfections were as follows: PSD-95-pTagRFP (Addgene, #52671), pmScarlet_C1 (Addgene, #85042), pEGFP-N1 (a gift from Pirta Hotulainen, University of Helsinki), and the CDKL5 cloned constructs GFP-CDKL5, GFP-CDKL5 L960D, or CDKL5-GFP.

### Imaging

Images were taken using a Marianas spinning disk confocal microscope (3i) using a Plan-Apochromat 100x/1.46 oil (Zeiss) objective and a FLASH4 sCMOS (Hamamatsu) camera.

### Primary cortical neuron imaging

Following 20-22 hr of transfections, DIV15 cortical neurons were fixed with 4% paraformaldehyde for 15 min at room temperature, washed 3 times with 1x phosphate-buffered saline (PBS) and mounted onto glass slides with ProLong® Diamond Antifade (ThermoFisher Scientific). Samples were imaged using Olympus FV4000 confocal microscope (at the Light Microscopy Unit, Institute of Biotechnology, supported by HiLIFE and Biocenter Finland) equipped with a UPLXAPO 60x/1.42 oil-immersion objective and SilVIR detectors. The images were acquired in 16-bit format with a pixel dimension of 0.1657x 0.1657. Images were further processed utilising Fiji ImageJ for analysis.

### Image Analysis

CDLK5 proximity to focal adhesions and mean nuclear to cytoplasmic ratio was calculated using ImageJ software. Adhesion masks were created by thresholding the vinculin focal adhesion marker. This mask was enlarged by 0.5 μm and applied to the CDKL5 image channel to measure the intensity of CDKL5 proximal to the vinculin focal adhesions. This intensity was compared to the total intensity of the CDKL5 construct to express the focal adhesion localisation as a percentage of the whole cell intensity. The nuclear to cytoplasmic ratio of each CDKL5 construct was calculated by dividing the mean nuclear intensity of the construct by the mean cytoplasmic intensity of CDKL5 from a sum projection of each cell. N = 5 independent experiments.

Quantification of CDKL5 and PSD-95 colocalization in primary neurons was performed utilizing the JACoP plugin in ImageJ (Bolte and Cordelières, 2006). The region of interest for CDKL5 and PSD-95 signals along the dendrites in the max intensity projections of Z-stacks were defined by manual thresholding for each sample to exclude the background region outside of dendritic shaft and spines. These images were further analysed with JACoP colocalization plugin to estimate Pearson’s correlation coefficient as a parameter for colocalization between CDKL5 and PSD-95.

For quantifying the spine density of neurons transfected with CDKL5 constructs, the mScarlet signal was used as a morphological marker to identify neuronal dendrites in the maximum intensity projections of the Z-stack. The number of spines along a given length of a dendritic shaft was counted and divided by the length of the adjacent shaft. The values were represented as relative dendritic protrusions per 10 µm length of the dendritic shaft, normalized to the control.

### Statistical analysis

To calculate and plot the means, standard deviation and statistical significance of the CDKL5 image analysis in cells and neurons, GraphPad Prism v.10.2.3 (GraphPad Software, San Diego, CA, USA) was used.

### Analysis of CDKL5 variants in ClinVar

The CDKL5 variant data used to generate Fig. 8 are from ClinVar with the following accession numbers;

RCV000413039.3, RCV004724834.1, RCV000169993.2, RCV000623570., RCV002318854.9, RCV000133318.4, RCV000560461.7, RCV000501064.6, RCV003765058.3, RCV001959030.7, RCV001089709.1, RCV000535244.9, RCV002249338.1, RCV000169916.4, RCV000133328.7, RCV004724846.1, RCV001253212.1, RCV000990486.1, RCV004724850.1, RCV001383236.3, RCV000133337.4, RCV004723156.1, RCV001942309.7, RCV000307290.1, RCV000548702.9, RCV001899898.5, RCV000145530.10, RCV000990488.1, RCV004724997.1, RCV004724736.2, RCV001172318.1, RCV000145533.16, RCV000144836.2, RCV000416943.1, RCV000144837.1, RCV000170034.2, RCV004724999.1, RCV000201936.1, RCV000803646.7, RCV000578897.5, RCV000699805.8, RCV002022681.7, RCV000622992.3, RCV000640488.8, RCV000170053.2, RCV004725011.1, RCV002312260.9, RCV001172319.1, RCV000133383.4, RCV001266418.3.

## Acknowledgements

B.T.G. was supported by Cancer Research UK Program Grant (CRUK-A21671) and British Heart Foundation Special Project Grant (SP/F/23/150045). V.S. and J.S. were supported by the Magnus Ehrnrooth Foundation, Research Council of Finland (363743), Sigrid Jus*é*lius Foundation (8029) and Jane and Aatos Erkko Foundation (240046). We acknowledge Kinga Grondkowska for early discussions and analysis and Dorus Gadella, Johannes Hell, and Pirta Hotulainen for sharing reagents.

## Supplementary Information

**Supplementary Figure 1.**
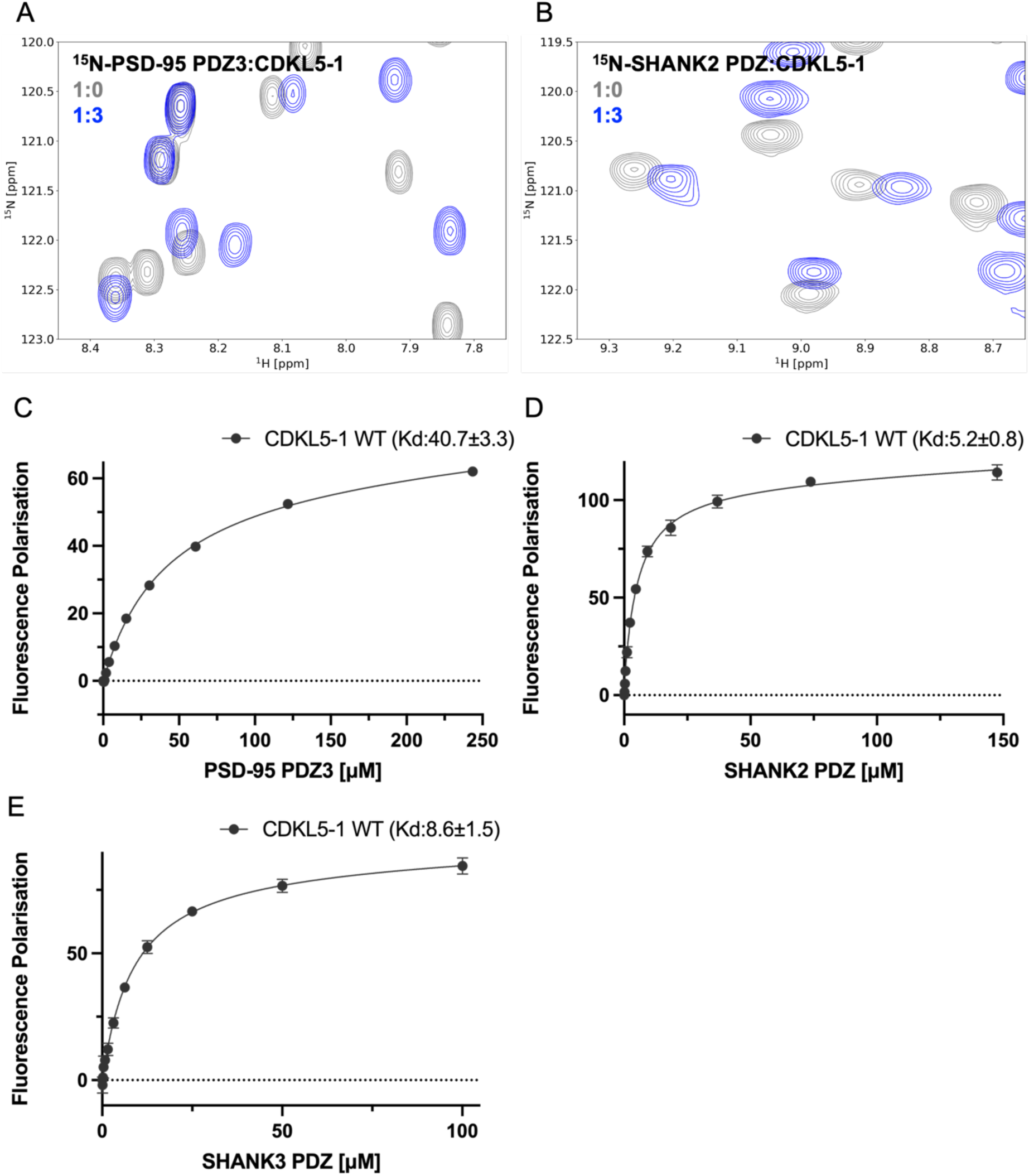
CDKL5 isoform 1 C-terminus binding to other PDZ domains. **(A)** NMR spectra of ^15^N-labelled PSD-95 PDZ3 with CDKL5-1^941-960^ peptide. PSD-95 PDZ3 on its own (grey), and with CDKL5-1^941-960^ peptide at 1: 3 (blue) molar ratio. **(B)** NMR spectra of ^15^N-labelled SHANK2 PDZ with CDKL5-1^941-960^ peptide. SHANK2 PDZ on its own (grey), and with CDKL5-1^941-960^ peptide at 1: 3 (blue) molar ratio. **(C-E)** FP assay for CDKL5-1^941-960^ peptide fluorescein-labelled via a non-native cysteine at the N-terminus against **(C)** PSD-95 PDZ3, **(D)** SHANK2 PDZ and **(E)** SHANK3 PDZ. Dissociation constants ± SE (µM) for the interactions are indicated in the legend. All measurements were performed in triplicate.

**Supplementary Figure 2.**
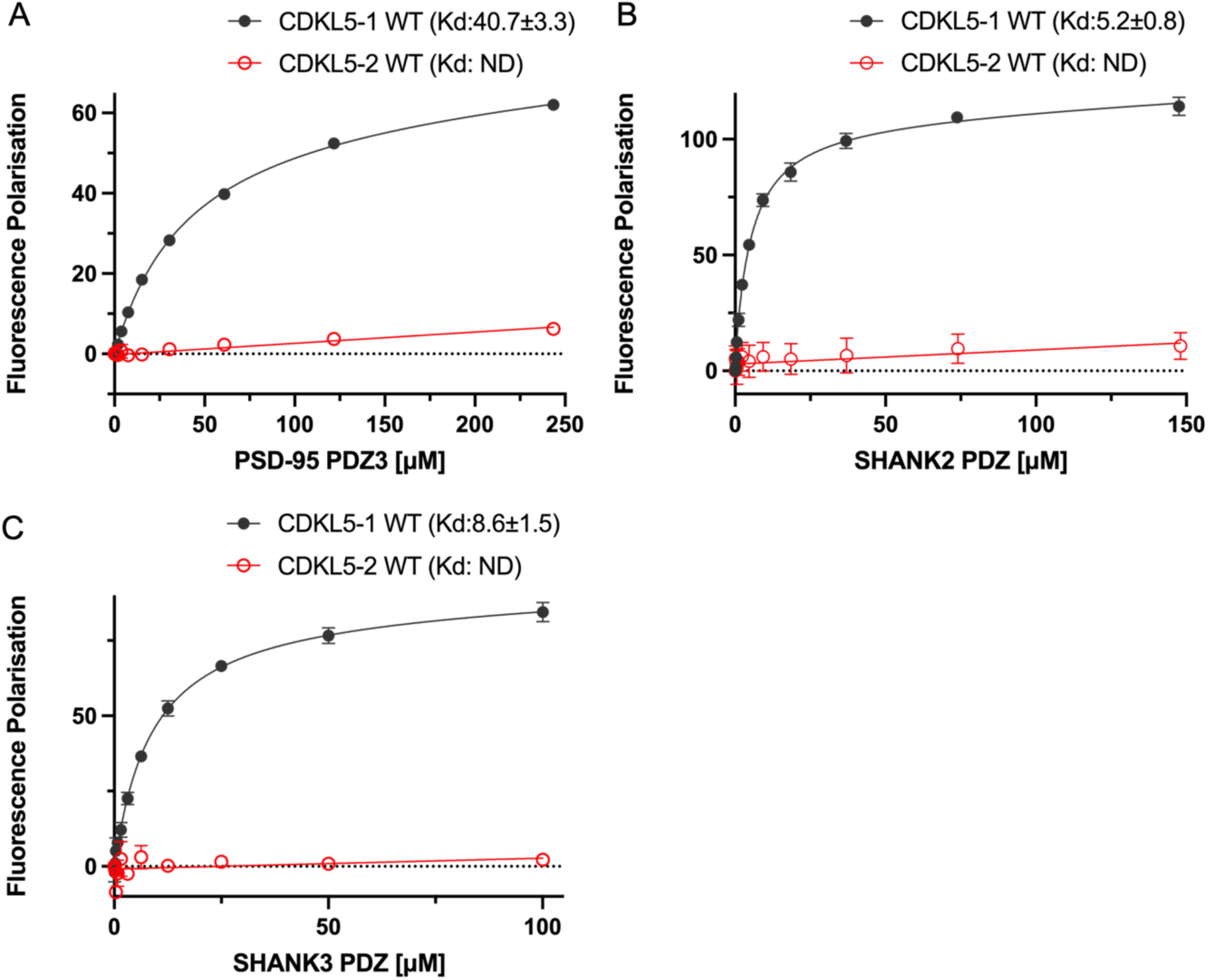
CDKL5 isoform 2 does not interact with PDZ domains. **(A-C)** FP assay for CDKL5-2^1011-1030^ peptide fluorescein-labelled via a non-native cysteine at the N-terminus against **(A)** PSD-95 PDZ3, **(B)** SHANK2 PDZ and **(C)** SHANK3 PDZ. Dissociation constants ± SE (µM) for the interactions are indicated in the legend. ND, not determined. In each panel the binding to CDKL5-1^941-960^ is shown. All measurements were performed in triplicate.

**Supplementary Figure 3.**
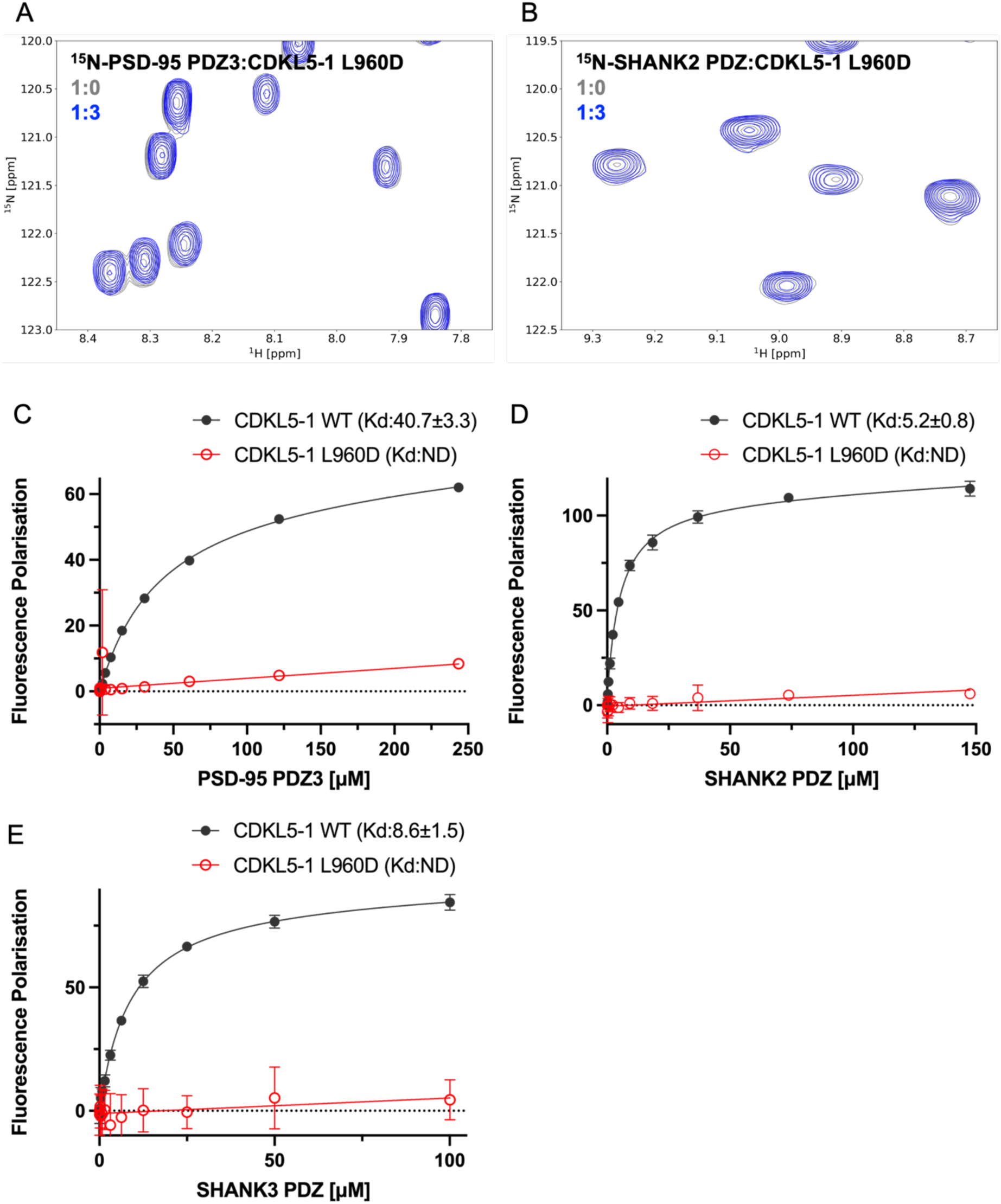
The L960D mutation in CDKL5 isoform 1 abolishes the interaction with PDZ domains. **(A)** NMR spectra of ^15^N-labelled PSD-95 PDZ3 with CDKL5-1^941-960^ L960D peptide. PSD-95 PDZ3 on its own (grey), and with CDKL5-1^941-960^ L960D peptide at 1: 3 (blue) molar ratio. **(B)** NMR spectra of ^15^N-labelled SHANK2 PDZ with CDKL5-1^941-960^ L960D peptide. SHANK2 PDZ on its own (grey), and with CDKL5-1^941-960^ L960D peptide at 1: 3 (blue) molar ratio. **(C-E)** FP assay for CDKL5-1^941-960^ L960D peptide fluorescein-labelled via a non-native cysteine at the N-terminus against **(C)** PSD-95 PDZ3, **(D)** SHANK2 PDZ or **(E)** SHANK3 PDZ. In each panel the binding to CDKL5-1^941-960^ is shown. All measurements were performed in triplicate. Dissociation constants ± SE (µM) for the interactions are indicated in the legend. ND, not determined.

**Supplementary Figure 4.**
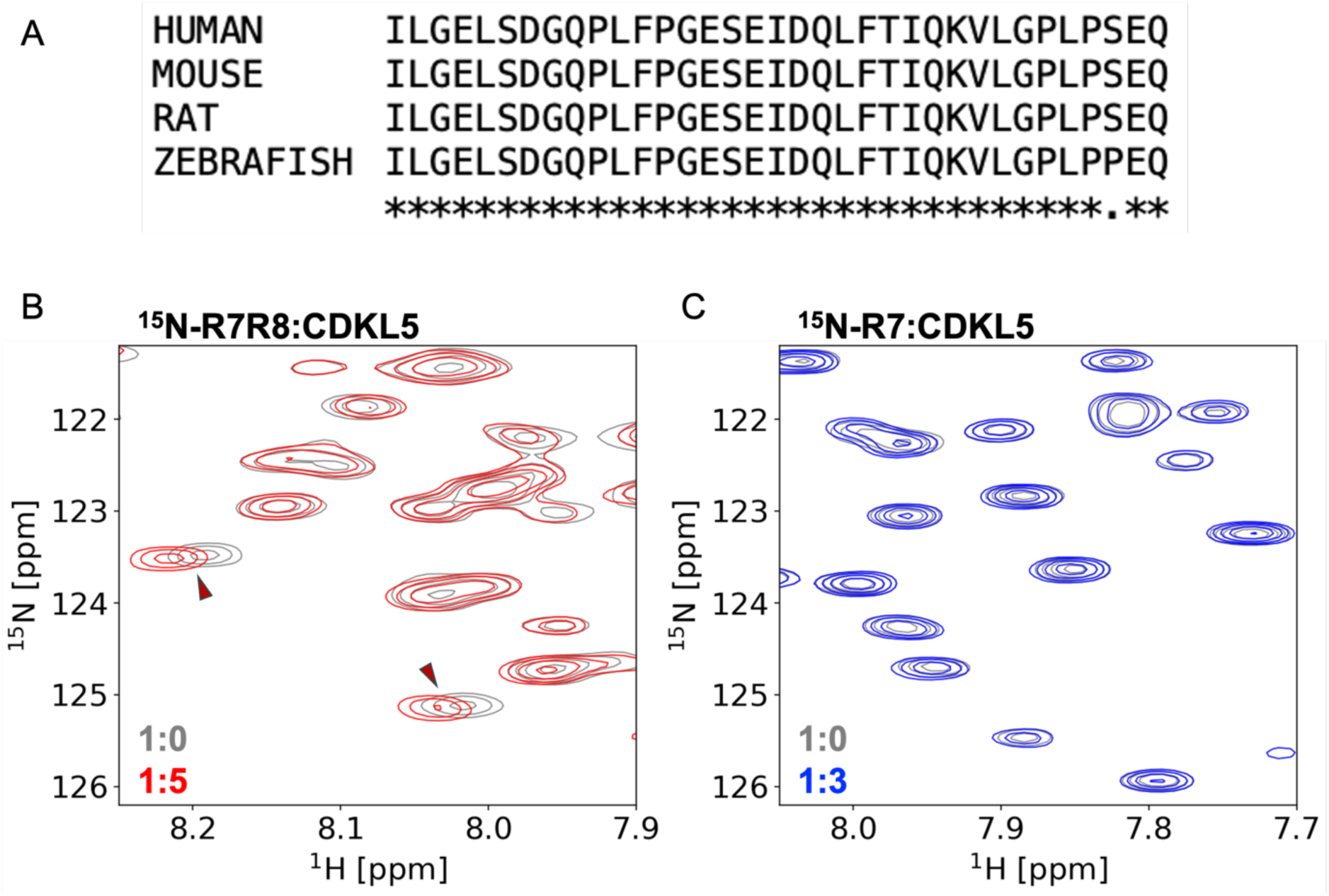
The CDKL5 kinase domain binds to talin1 R7R8 but not to R7. **(A)** Sequence alignment of the talin-binding site region in CDKL5 from human, mouse, rat and zebrafish (UniProt: B9X2H5, O76039, Q3UTQ8, E2E1S0). **(B-C)** NMR spectra of ^15^N-labelled talin1 R7R8/R7 with CDKL5-1^213-234^ peptide. **(B)** talin1 R7R8 on its own (grey) and with CDKL5-1^213-234^ peptide at 1: 3 (red) molar ratio. **(C)** talin1 R7 on its own (grey), and with CDKL5-1^213-234^ peptide at 1: 3 (blue) molar ratio.

**Supplementary Figure 5.**
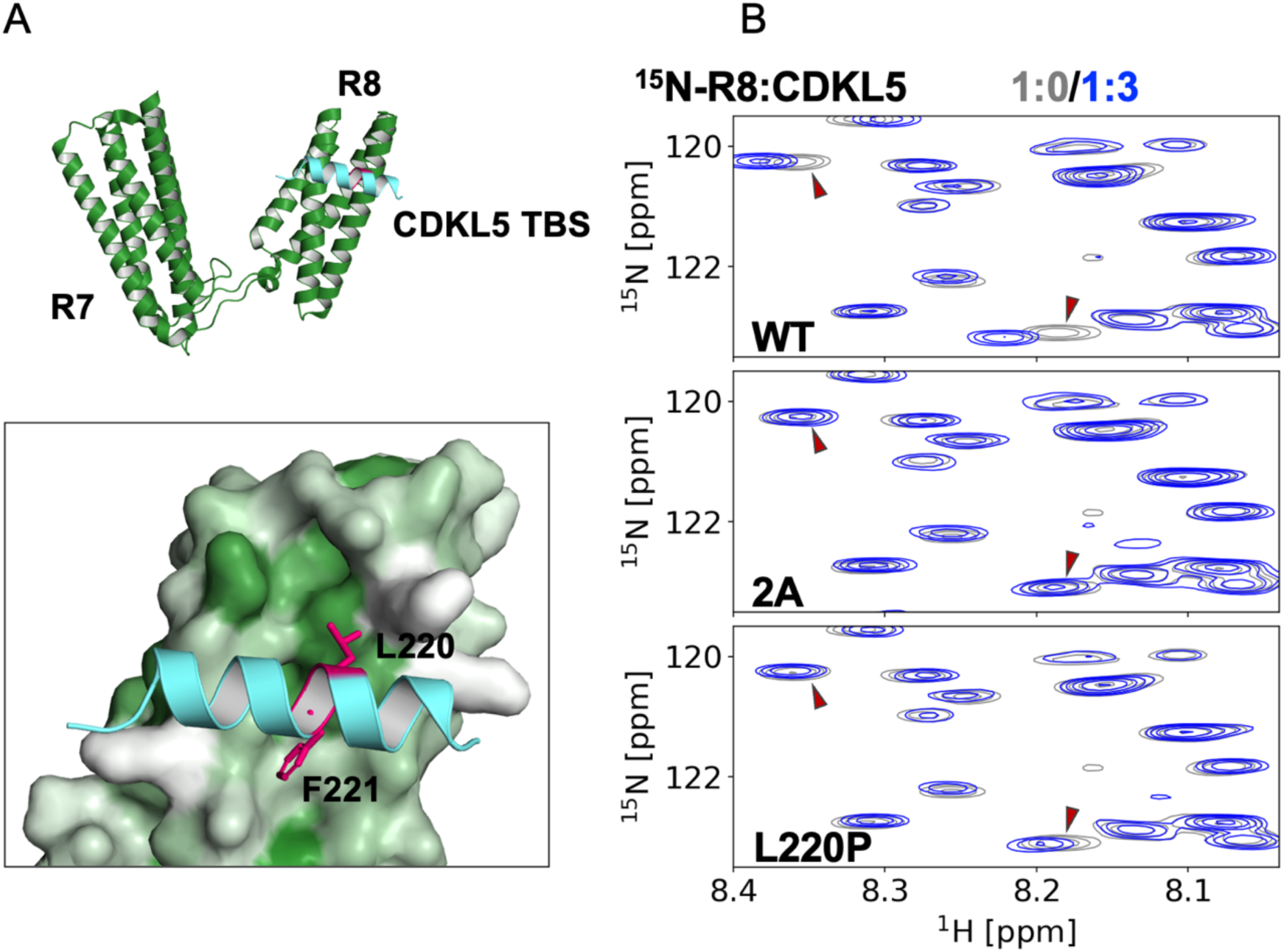
Mutations in CDKL5 to disrupt binding to talin R8. **(A)** Structural model of R7R8 (green) in complex with the CDKL5 talin-binding motif peptide (cyan) based on the R7R8:CDK1 crystal structure (PDB: 6TWN) (inset) the binding surface between R8 and CDKL5 showing the two mutated residues L220 and F221 (pink). The talin R8 surface is coloured by hydrophobicity using the AA index database (entry FASG890101 [Nakai et al., 1988]) where hydrophobic residues are shown in green and polar residues in white. **(B)** NMR spectra of ^15^N-labelled talin1 R8 on its own (grey), and with CDKL5^213-234^ WT (top), CDKL5^213-234^ 2A (middle) and CDKL5^213-234^ L220P (bottom) peptides at a 1:3 ratio (blue). Red arrows indicate peaks shifted in the presence of wildtype CDKL5^213-234^ peptide.

**Supplementary Figure 6.**
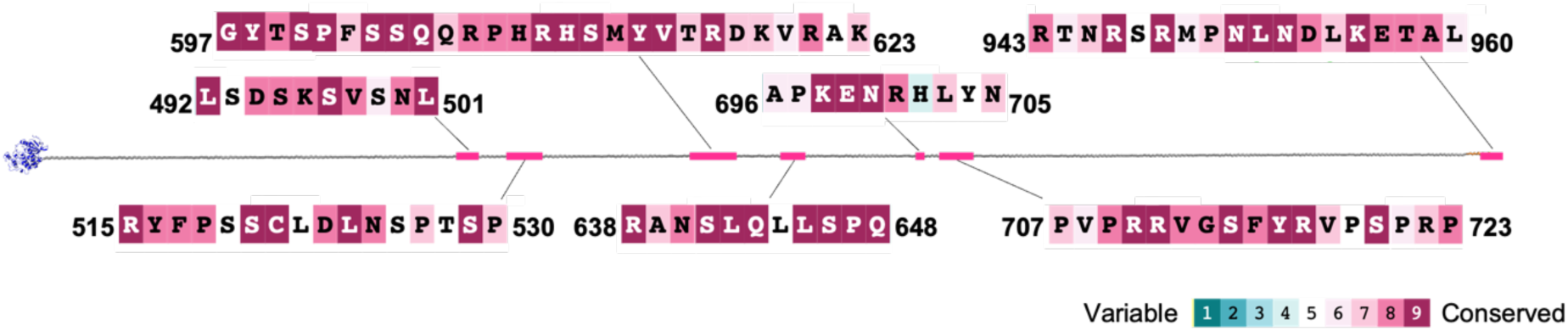
The CDKL5 unstructured region contains seven highly conserved regions. Centre, the unfurled CDKL5 structure as in Fig. 1D, the kinase domain is shown in blue. Seven regions containing more than 4 residues in a row with conservation score 8 or 9 are highlighted with magenta. The very C-terminal highly conserved region is the PDZ-binding motif. The primary sequences of the seven highly conserved peptides are shown. Residues are coloured based on the ConSurf conservation score (Yariv et al., 2023) where magenta denotes high conservation.

